# Exploring the role of EMT in Ovarian Cancer Progression: Insights from a multiscale mathematical model

**DOI:** 10.1101/2024.06.25.600568

**Authors:** Samuel Oliver, Michael Williams, Mohit Kumar Jolly, Deyarina Gonzalez, Gibin Powathil

## Abstract

Epithelial-to-mesenchymal transition (EMT) plays a key role in the progression of cancer tumours and can make treatment significantly less successful for patients. EMT occurs when a cell gains a different phenotype and possesses different behaviours to those previously exhibited. This may result in enhanced drug resistance, higher cell plasticity, and increased metastatic abilities. It has therefore has become essential to encapsulate this change and study tumour progression and its response to treatments. Here, we use a 3D agent-based multiscale modelling framework based on Physicell to investigate the role of EMT over time in two cell lines, OVCAR-3 and SKOV-3. The impact of conditions in the microenvironment are incorporated into the model by modifying cellular behaviours dependant on variables such as substrate concentrations and proximity to neighbouring cells. OVCAR-3 and SKOV-3 cell lines possess highly contrasting tumour layouts, allowing a vast array of different tumour dynamics and morphologies to be tested and studied. The model encapsulates the biological observations and trends seen in tumour growth and development, thus can help to obtain further insights into OVCAR-3 and SKOV-3 cell line dynamics. Sensitivity analysis was performed to investigate the impact of parameter sensitivity on model outcome. Sensitivity analysis showed that parameters used in generating the rate of EMT and cycle rates within the cells are relatively more sensitive than other parameters used.

## 1 Introduction

Epithelial-to-Mesenchymal Transition (EMT) is a process in which epithelial cells undergo phenotypic changes, enabling a reduction in cell-cell adhesion and enhancing migratory abilities [1, 2]. EMT is essential for normal tissue functionality within the body [3]. It allows the closure of developmental neural tubes [4], plays a key involvement in embryogenesis [5], and enables wound healing to occur [6]. Despite the reliance of the human body on this process, the role of EMT can occasionally become detrimental and further complicate treatment for diseases. EMT is heavily linked to cystic fibrosis by causing goblet cell and pneumocyte hyperplasia in the lungs [7]. Rheumatic diseases have also been linked to EMT, with rheumatoid joints expressing an abundant amount of TGF-*β* in the synovial fluid [8, 9].

It was recently suggested to separate EMT into three main types [10, 11]. EMT occurring during a self-contained process requiring multiple cell types to be generated such as organ development and embryo formation is classified as type 1. EMT associated with repair such as wound healing, organ fibrosis, and tissue regeneration is classified as type 2. This repair discontinues upon completion and when inflammation is reduced. Type 3 EMT includes instances where there is a genetic and epigenetic difference between the epithelial and mesenchymal cell types. This is the key type of EMT associated with cancer progression and metastasis.

EMT is no longer considered a binary switch [12, 13]. Instead, cells can fluctuate through a multi-step process during which they may show partial epithelial and mesenchymal characteristics [14]. This leads to a more complex differentiation process between classifications of cells. Various biological markers are used to conclude the placement of these cells along the EMT scale [15]. E-Cadherin is a surface marker used for the identification of epithelial cells [16], while N-Cadherin, vimentin, and fibronectin are markers to locate mesenchymal cells [17]. The ratio of these markers can be used to conclude the epithelial/mesenchymal balance determined for each individual cell [18].

EMT is a crucial step in the cancer progression [19]. Mesenchymal cells within a tumour have lower cell-cell adhesion forces due to a reduced E-Cadherin expression on the cell surface [20]. This allows the cells to break away from the main tumour location and escape from the brick-like structure they were previously a part of [21]. This relocation of cells can cause metastasis away from the primary tumour site. Metastatic cases are responsible for over 90% of all cancer-related deaths [22]. One justification for this is the improved drug resistance possessed by the slower cycling metastatic cells [23, 24]. These cells obtain resistance to anoikis and switch from a phenotype tailored for proliferation to a phenotype which targets invasion around the body [25]. This lack of proliferation hinders the effectiveness of the drug, as targeting the rapidly dividing cells is no longer efficacious [26, 27]. This effect is responsible for lower long-term treatment dosages occasionally being beneficial. Higher dosages can eradicate the susceptible, less concerning epithelial cells, therefore making space and freeing up resources for the mesenchymal cells to exploit [28, 29, 30]. This paper will focus on the link between the tumour microenvironment and the initialisation of EMT within cells [31, 32].

It has been shown that cells are also capable of undergoing mesenchymal-to-epithelial transition (MET) [33, 34]. This is the reverse process of EMT, where mesenchymal cells transition back to epithelial cells and regain the epithelial phenotype and behaviours previously exhibited [35, 36, 37]. The phenomenon of MET is primarily seen in relocated mesenchymal cells which return to focusing on proliferation [38]. While completing EMT allows a cell to travel with more ease throughout the body, MET enables transformation back to the faster proliferating, more stable epithelial phenotype [39, 40]. This allows the tumour to grow faster and spread throughout the body more rapidly [41, 42]. With a higher focus on the role of EMT in recent years [43, 44], mathematical models are becoming a valuable asset to scientific research into improving survival and cancer treatment success rates [45, 46, 47]. *In-silico* models offer a fast and accurate alternative to *in-vivo* or *in-vitro* models [48], both of which expend physical resources such as cell lines to perform the experiments, require time for the biological processes to complete, and need potentially complex ethical and moral justifications [49, 50]. These models can be used to test the impacts of different parameter values or conditions in the microenvironment by using simple adaptations, eventually informing experimental settings [51].

Here, we investigate the role of EMT in the progression of ovarian cancer by developing a multiscale mathematical and computational model to investigate the role of EMT in cancer cells with a focus on two common ovarian cancer cell lines: OVCAR-3, a human ovarian epithelial carcinoma cell line [52] and SKOV-3, a human ovarian adenocarcinoma cell line [53]. This model allows cells to migrate through the domain, proliferate at microenvironment-dependent rates, and progress through EMT in a biologically realistic manner. The model is based on experimental data and processes found in past literature. During the simulations, information on each cell such as position, velocity, surrounding oxygen levels, and cell cycle stage can be tracked and analysed.

The model will also be used to study the importance of including MET in tumours, a key process when modelling metastatic cancers. Direct comparisons will be made between simulations both including and excluding the presence of MET. Sensitivity analysis on the key parameters is performed to help quantify the role of EMT and its association with tumour size and composition over time. Following model validation using experimental data, future predictions are made on different initial states of the tumour. Temporal dynamics of cells initialised with epithelial and mesenchymal cells across different cell lines are compared, with the importance of accurately distinguishing the cell lines highlighted.

## 2 Mathematical Model and Methods

The model developed here is built upon a Physicell framework [54, 55]. Physicell is an agent-based, multiscale, 3D framework used to perform *in-silico* simulations of biological processes. Cells move off-lattice along a continuous domain. Substrates are tracked on a 3D discrete mesh into which the domain is divided. A detailed overview of the cell and substrate mechanics can be found in the Physicell overview paper by Macklin et al [54, 55].

### 2.1 Cadherin Rating

The two cell lines used here to investigate the EMT process possess highly contrasting characteristics, making the mathematical model more versatile and adaptable to cell lines. Figure 1 shows the cross-section of (a) OVCAR-3 and (b) SKOV-3 tumours in an *in-vitro* experiment. SKOV-3 cells are seen to show a high expression of N-Cadherin, suggesting a significant mesenchymal cell population within the tumour [56]. During biological observations of SKOV-3 spheroid experiments, a thin shell of epithelial cells located around the periphery of the tumour is created. OVCAR-3 cells are seen to possess more epithelial behaviours, expressing high amounts of the marker E-Cadherin [56]. While the majority of OVCAR-3 cells in the spheroid are epithelial, small clumps of mesenchymal cells are formed inside the tumour. This creates a polka dot effect in the observations of the neoplasm.

**Figure 1:**
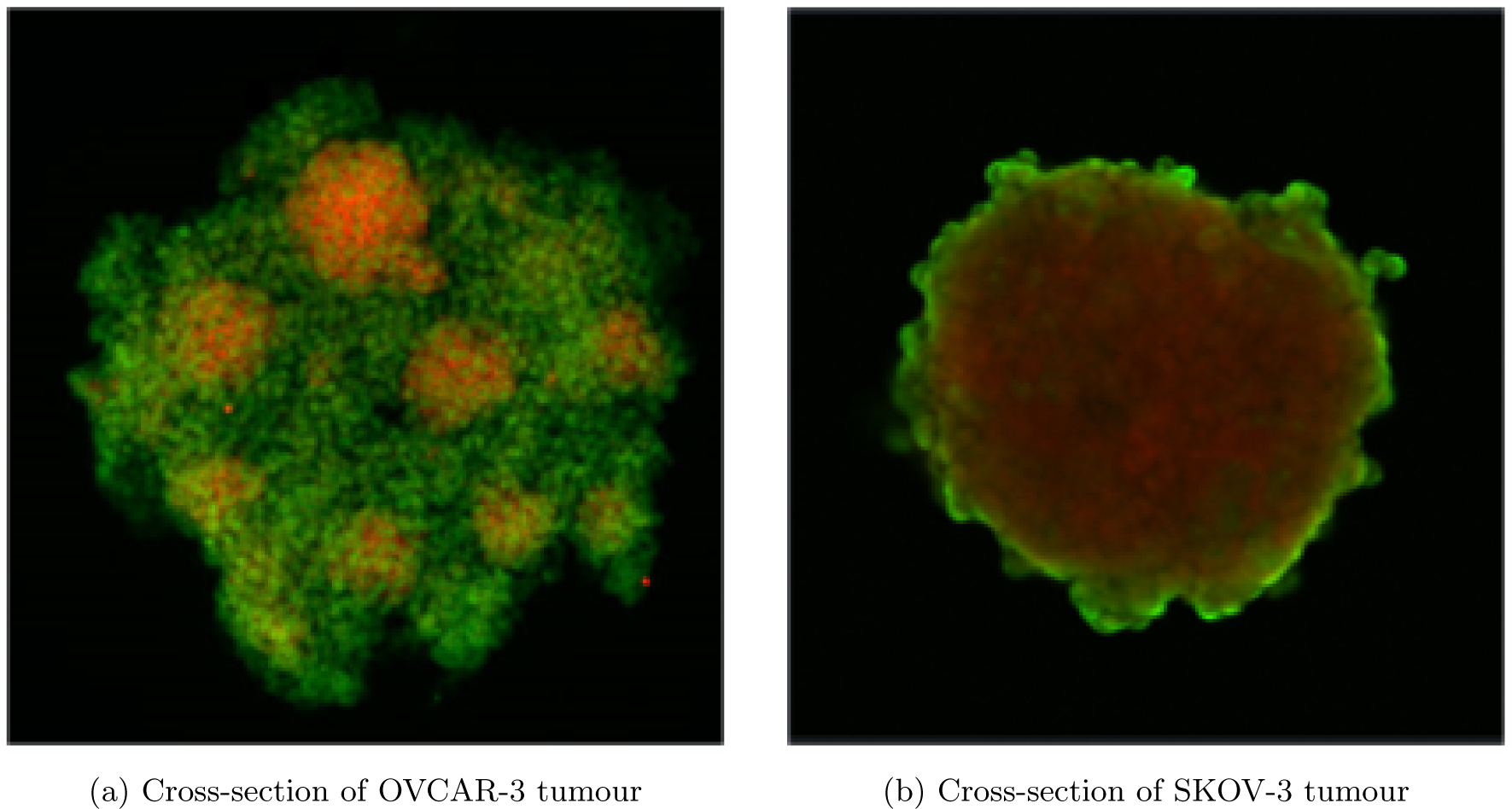
Cross-section of tumours upon experiment completion (unpublished). Green areas represent high levels of E-Cadherin, expressed in epithelial cells. Red areas show high expression of N-Cadherin, a marker for mesenchymal cells. Small clumps of mesenchymal cells are shown in OVCAR-3 tumours surrounded by a backdrop of epithelial cells (a). Mesenchymal cells make up the majority of SKOV-3 tumours, with a thin layer of epithelial cells appearing on the periphery (b).

In the modelling framework, cells are assigned a rating along a scale of their epithelial/ mesenchymal phenotype running from 0 (epithelial) to 13 (mesenchymal), as illustrated in Figure 2. The maximum rating is set to 13 to allow an equally weighted assignment of epithelial cells to ratings of 0-6 and mesenchymal to 7-13. Moreover, a central rating value for epithelial (3) and mesenchymal (10) cells can be set as the default if required. On each iteration, a cell has a probability of increasing cadherin rating depending on various inter-cellular and intra-cellular conditions such as hypoxia and current cell rating.

**Figure 2:**
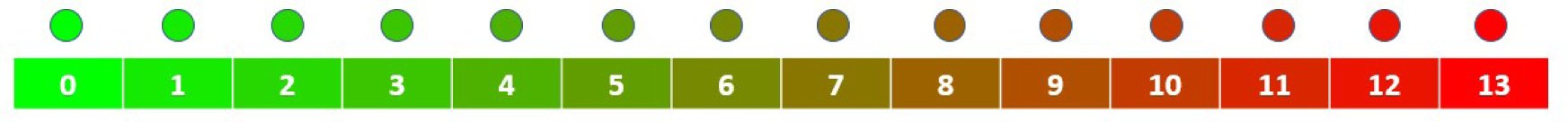
Appearance of cells during the simulations based on their current cadherin rating.

To investigate EMT, cells are initiated with a cadherin rating of zero. It is assumed that only EMT can occur since we take the tumour to be in its primary location [57]. MET, the reverse process, primarily only occurs when the mesenchymal cells have relocated and stabilised in a new environment [58, 59]. This MET process will be discussed in more detail in Section 4. When a parent cell divides, the cadherin rating in the model is conserved to the daughter cells.

The cadherin rating of a cell has a key influence on its phenotypic characteristics [60]. We use a number of Hill functions to build correlations between the current cadherin rating and the behaviour of a cell. Table 1 shows the impact that higher values of the cadherin rating have on the cell parameters used in the model. Here, we adopt other cellular parameter values from the Physicell model framework [55, 54].

**Table 1:** Impact of increase in cadherin rating on various cell behaviours.

| Cadherin Rating Impact | Behaviour | Figure 3 |
| --- | --- | --- |
| Increases | Migration Speed | (a) |
| Decreases | Cell-Cell Adhesion | (b) |
| Increases | Signal Secretion | (c) |

Figure 3 shows the assumed quantitative trends with cadherin rating changes in different cell behaviours, such as migration speed (a), adhesion strength (b), and signal secretion rate (c). Mesenchymal cells have been found to possess enhanced migratory tendencies [61, 62] and lower adhesion strength than seen in epithelial cells [63, 64]. Here, the increased migration speed which is permitted for mesenchymal cells allows the cells to move with greater freedom through the domain and increase the probability of the cell leaving the tumour itself to reach the domain boundary. The decrease in adhesion strength allows the mesenchymal cells to break away from neighbouring cells and escape the tumour with greater ease. Signal secretion rate is increased with cadherin rating, meaning epithelial cells around mesenchymal cells are in a higher concentration of signal and are more likely to increase in cadherin rating themselves, inducing the bystander effect.

**Figure 3:**
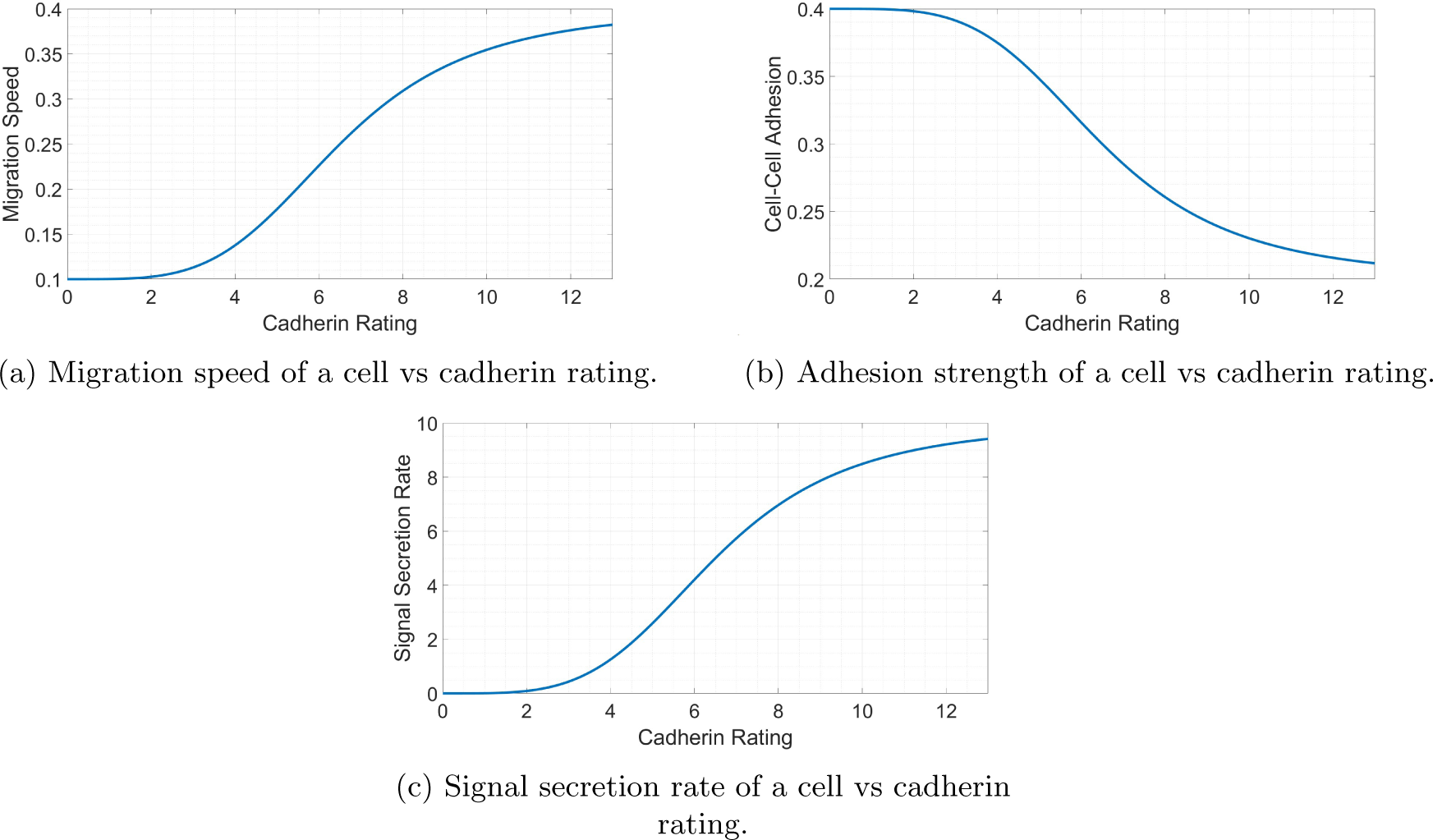
Dependence of different cell variables on the current cadherin rating of the cell. Increased cadherin ratings increase cell migration and bystander signal secretion rate while decreasing cell-cell adhesion strength.

Here, by bystander effects, we mean the cumulative effects of various factors such as cytokines, growth factors, and exosomes secreted by cancer cells [65]. In the model, the bystander effect is responsible for the formation of the disjoint mesenchymal clumps observed in the OVCAR-3 hybrid tumour population, co-expressing epithelial and mesenchymal markers [66]. We assume that these bystander effects encourage epithelial cells in their proximity to undergo EMT. These factors secreted by mesenchymal cells help encourage changes in the cell phenotype and increases the rate at which cells undergo EMT [67, 68]. This creates localised pockets of mesenchymal clusters in which high amounts of these secreted factors are present, further encouraging EMT in the surrounding cells. Such tumour heterogeneity is apparent in solid tumours where mosaic expression of cadherins have been described in both primary and secondary epithelial ovarian cancer tumours [69, 70]. Thus our model recapitulates phenotypic heterogeneity and mechanisms of chemoresistance which are apparent in solid tumours and in tumour spheroids by incorporating aspects of a localised tumour microenvironment including gradients of nutrient availability and cellular necrosis [71].

### 2.2 Cell Cycle

Cells in the model progress through the cell cycle at varying rates depending on the conditions within both the cell and the microenvironment (See Figure 4) [72]. Cells with a higher cadherin rating are assumed to have more mesenchymal phenotypic behaviours and so cycle slower than those possessing more epithelial characteristics [73]. We include this phenomenon by introducing a Hill function used in generating a cadherin cycling impact parameter, as shown in Figure 4 (a). These Hill functions are given a Hill power of four. This value is large enough to allow small changes in the behaviours when for small changes in dependant variables, while being large enough to create non-linear correlations. This parameter ranges from two for entirely epithelial cells to just above one for entirely mesenchymal cells. Cells under higher pressure due to combined repulsion forces from neighbouring cells also reduce the cell cycling rate, as shown in Figure 4 (b) [74, 75]. This is to encapsulate the effect of cells requiring empty space in the surrounding area to divide into. We generate this parameter value using a similar Hill function to that seen in Figure 4 (a), where the parameter value ranges between two where no pressure is applied to the cell and just above one where the pressure is two units. With a greater concentration of oxygen available in the microenvironment cells are able to increase adenosine triphosphate (ATP) production and therefore cycle faster [76, 77, 78, 79]. This is incorporated into the model in Figure 4 (c) by introducing an oxygen impact parameter ranging from one in hypoxic conditions to nearly two in oxygen-rich conditions.

**Figure 4:**
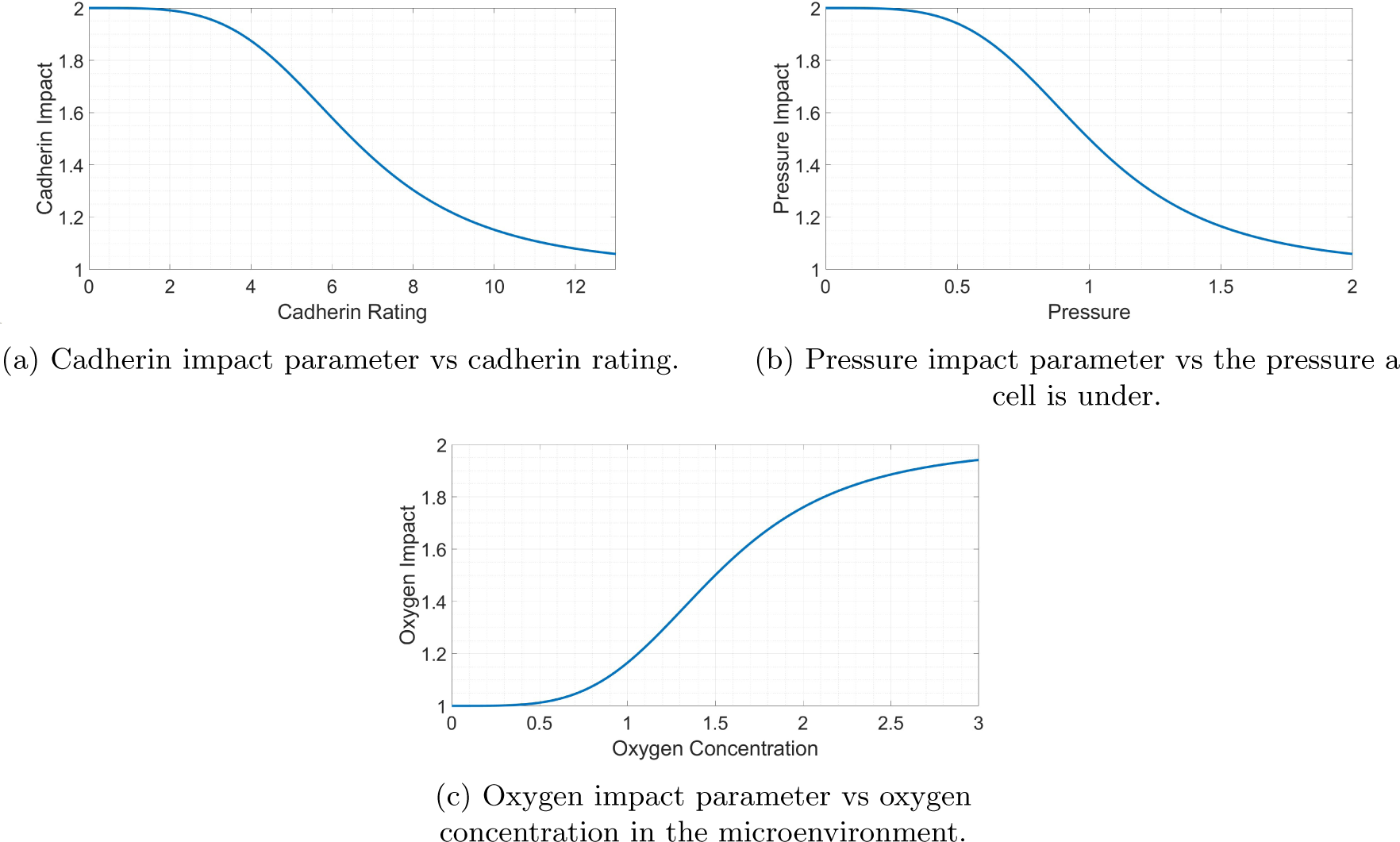
Dependence of cell cycling parameter values on cell conditions.

These parameters are used to calculate the rate at which a cell leaves the G1 stage of the cell cycle using Equation 1, where *r* is the cycling rate, *n* is the current population, and *K* is the carrying capacity. The carrying capacity is set to 6500 to allow a maximum population similar to that observed in the experimental data. Due to the cycling rate variability being enforced in the G1 stage of the cycle, there is a time delay in the effect of this rate and the tumour can reach populations higher than that specified in the carrying capacity of the logistic growth term. We include the logistic growth term with a carrying capacity to reduce the growth rate of large tumours, observed experimentally in which populations plateau.

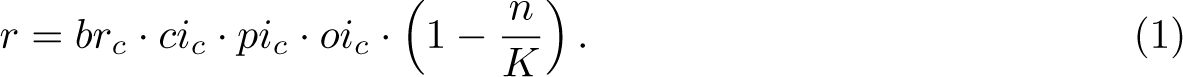

We denote *br_c_*as the base cycle rate, *ci_c_*as the cadherin impact on the cycling rate, *pi_c_* as the pressure impact on the cycling rate, and *oi_c_*as the oxygen impact on the cycling rate. Assuming average cells leave G1 with a rate of 1/11 hours*^−^*^1^ [80], these parameters allow a maximum cycling rate of 8/11 hours*^−^*^1^ to leave G1. The length of the cell cycle would therefore vary between around 14 hours in the optimal conditions for cell proliferation and 24 hours in the poorest. The remaining steps in the cell cycle are unaffected by the microenvironmental conditions in the model.

### 2.3 Jump Probability - Incorporation of EMT

Here, we assume that not all cells have the same predicted rate of moving up through the cadherin rating along the EMT scale. On each iteration of the simulation, a cell is given a probability of moving up through the cadherin rating and gaining more mesenchymal properties. We assume that the magnitude of this probability is dependent on factors in both the cell itself and the microenvironment around it. These inter-cellular and intra-cellular conditions give rise to a stochastic process by which the cadherin rating of the cell is determined.

#### 2.3.1 Impact of Inter-Cellular Conditions

Following experimental observations, we hypothesise in our model that two main factors in the microenvironment contribute to EMT within epithelial cells. Hypoxic conditions have been found to encourage EMT in cancer cells by generating various signalling pathways and activating transforming growth factor TGF-*β* [81, 82]. This is achieved in the model using a Hill function to produce an oxygen impact parameter decreasing from one in hypoxic conditions to approximately zero in oxygen-rich microenvironments, as shown in Figure 5 (a). It has been observed experimentally that mesenchymal cells appear to promote EMT [83]. Here, we incorporate this using “bystander signals” and is modelled with a PDE. Cells higher on the cadherin rating scale secrete the chemical signal responsible for the bystander effect at increased rates. This leads to higher signal concentrations around mesenchymal cells’ location due to the low diffusion coefficient (0.1 microns^2^/min). This creates a chain reaction of EMT within the cells and is responsible for the clumps observed in the OVCAR-3 spheroids seen throughout the *in-vitro* experiments. Figure 5 (b) shows the quantitative impact the signal concentration in the microenvironment has on the signal impact parameter.

**Figure 5:**
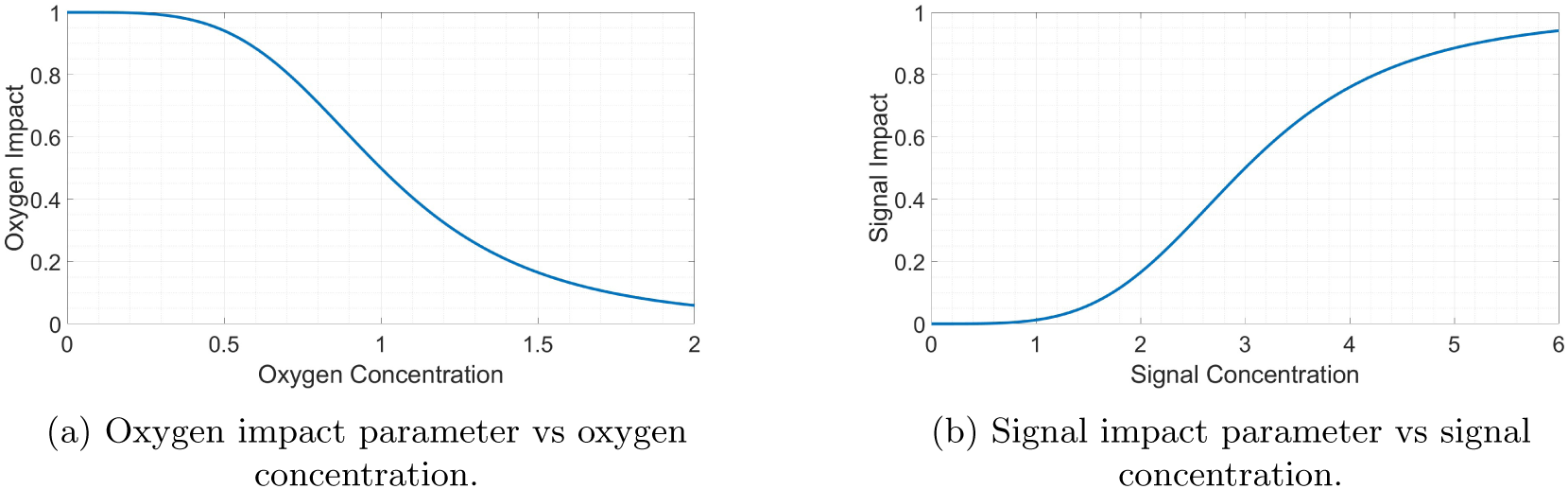
Dependence of the EMT rate parameters on the microenvironment.

#### 2.3.2 Impact of Intra-Cellular Conditions

Cells are also given a “base EMT probability” variable, causing cells with a higher cadherin rating to progress faster through the EMT scale. This has been observed biologically, in which epithelial and mesenchymal cells possess the most stability, with hybrid cells considered metastable with the highest plasticity [84, 85]. We introduce this phenomenon into the model using a Hill function based on the current rating of a cell, as illustrated in Figure 6. The parameter ranges from zero for purely epithelial cells to near one for purely mesenchymal. We combine this variable with the “oxygen EMT impact” and “signal EMT impact” described in Section 2.3.1.

**Figure 6:**
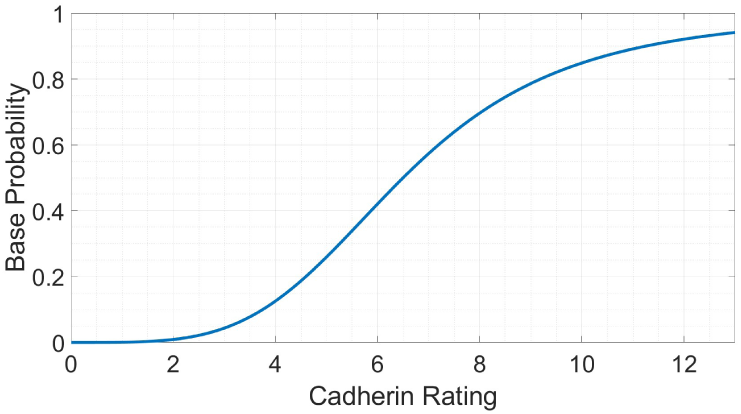
Base probability parameter vs current cadherin rating.

#### 2.3.3 Cumulative Effect on Jump Probability

The variables discussed previously in Section 2.3 are incorporated into an overarching equation, Equation 2, for OVCAR-3 and SKOV-3 cells, showing their cumulative effect. This “jump probability”, *p*, determines if a cell will increase its cadherin rating during each six-minute iteration of the computer simulation, leading to slightly increased mesenchymal behaviours highlighted in Section 2.1. The parameters in Equation 2 are cell line dependant, the details of which are highlighted in the Sections 2.3.4 and 2.3.5.

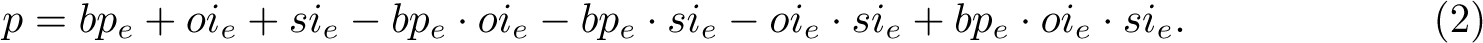

#### 2.3.4 OVCAR-3

We denote *bp^∗^* as the base EMT probability, *oi^∗^* the oxygen impact, and *si^∗^* the signal impact on rates of EMT in cancer cells shown in Figures 5 and 6. A weighting term is incorporated into these impact parameters, for which the value of each is shown in Table 2. The base EMT impact parameter has a low weighting to ensure the number of clumps arising in the tumour throughout the 4-day simulation is not unrealistically high (See Figure 1). The impact of oxygen is medium, as hypoxia is not seen as a requirement for EMT but does act as a key catalyst for the process [57, 86]. The chemical signal impact responsible for the bystander effect has a large weighting to ensure disjoint clumps can be formed quickly despite the low diffusion of the signal (see Figure 1). We assign an “EMT impact factor” parameter to both cell lines. This parameter quantifies the tendency for the cells to undergo EMT on each iteration of the simulation, depending on whether the cells are taken from OVCAR-3 or SKOV-3 cell lines. Since EMT does not occur in the majority of OVCAR-3 cells during experimental observations, OVCAR-3 is given a low impact factor of *α* = 1. Larger values of *α* lead to larger weightings of the parameters used in Tables 2 and 3 when generating the jump probability term, *p*, in Equation 2. We analyse the impact of the value of this *α* parameter in the Supplementary Material (SM).

**Table 2:** Values of the different variables used within the EMT calculation in Equation 2 for OVCAR-3 cells. The signal impact parameter has a large weighting to ensure sufficient levels of signal concentration can induce mesenchymal clump formation. The base EMT probability parameter has a small weighting to ensure EMT does not occur too frequently within the tumour leading to a scenario in which the mesenchymal clumps begin to connect.

| Parameter Name | Unweighted Parameter Symbol | Parameter Weight | Weighted Parameter Symbol |
| --- | --- | --- | --- |
| base EMT probability | $bp_e^*$ | 0.001 | $bp_e = 0.001 \cdot bp_e^*$ |
| oxygen EMT impact | $oi_e^*$ | 0.002 | $oi_e = 0.002 \cdot oi_e^*$ |
| signal EMT impact | $si_e^*$ | 0.01 | $si_e = 0.01 \cdot si_e^*$ |

**Table 3:** Values of the different variables used within the EMT calculation in Equation 2 for SKOV-3 cells. All parameters have an increased weighting to those used for OVCAR-3 cells to ensure sufficient

| Parameter Name | Unweighted Parameter Symbol | Parameter Weight | Weighted Parameter Symbol |
| --- | --- | --- | --- |
| base EMT probability | $bp_e^*$ | 0.005 | $bp_e = 0.005 \cdot bp_e^*$ |
| oxygen EMT impact | $oi_e^*$ | 0.01 | $oi_e = 0.01 \cdot oi_e^*$ |
| signal EMT impact | $si_e^*$ | 0.05 | $si_e = 0.05 \cdot si_e^*$ |

#### 2.3.5 SKOV-3

Unlike OVCAR-3 tumours, in SKOV-3 spheroids the red mesenchymal clumps are no longer distinguishable and instead a large pool covering the entire center of the SKOV-3 tumour is formed, as shown in Figure 1 (b). To ensure sufficient amounts of EMT occur to encapsulate this effect, the jump probability weightings are increased by a factor of five to that implemented for OVCAR-3 cells, as shown in Table 3. The jump probability is generated as shown in Equation 2 to ensure the biological observations remain in agreement with the simulation outputs. The shell of epithelial cells around the exterior of the tumour, as seen in experiments, is implemented by including an oxygen-dependent condition on the SKOV-3 cells. Hypoxic conditions in SKOV-3 tumours have been shown to upregulate the chemokine receptor CCR7, in turn inducing EMT development [87]. When oxygen concentration increases above a threshold value (set to four units), it is assumed mesenchymal cells undergo instant MET and are assigned the cadherin rating value of zero. This occurs only around the exterior of the tumour where oxygen is sufficient enough to cross this threshold. Table 3 shows the weightings of each parameter involved in generating the EMT probability for SKOV-3 cells. These increased weightings compared to those in Table 2 lead to vastly increased amounts of EMT when compared to OVCAR-3 tumours. The “EMT impact factor” parameter is given a value of five for SKOV-3 cells, meaning the parameter weights are five times larger in Table 3 than in Table 2. This value ensures sufficient EMT occurs throughout the SKOV-3 tumour to allow the pool of interior mesenchymal cells to develop inside the tumour.

### 2.4 Inititalisation

Here, we initialise the simulations with 1027 cells in a 3D spheroid scattered randomly within a sphere of radius 120 microns. This is to recreate the initial conditions as closely as possible to those used experimentally. Both oxygen and bystander signal substrates are absent at initialisation. Oxygen has a constant influx into the system through Dirichlet boundary conditions applied on all boundaries of the domain set to 38 units. The signal substrate is given Neumann boundary conditions with no constant influx. The model is simulated for 96 hours in which all cells are initialised with an equal cadherin rating, specified for each simulation.

An example of the initialisation is given in Figure 7. This depicts a cross-section of an OVCAR-3 tumour through the *z* = 0 plane at time *t* = 0 in which all cells are epithelial with a cadherin rating of zero.

**Figure 7:**
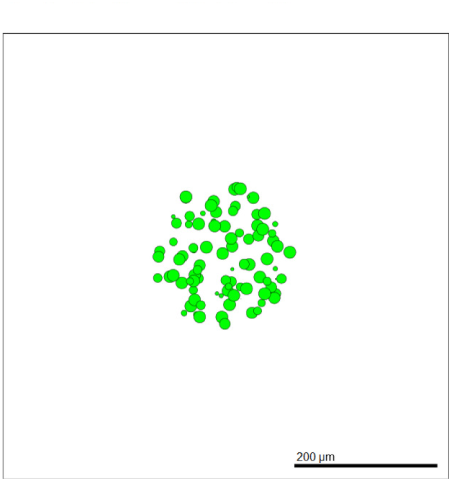
Cross-section of an example tumour at time *t* = 0.

## 3 Results and Discussions

Here, we study temporal tumour evolution over 96 hours of simulated time. OVCAR-3 and SKOV-3 epithelial cells are placed in the domain, with the cross-sections of the tumour shown each day until the completion of day four. The simulation results are then compared to the biological observations and data obtained from the *in-vitro* experiments.

### 3.1 OVCAR-3

After simulation completion, we can investigate how the tumours change in both composition and size over time. Figure 8 shows simulated cross-sections of an OVCAR-3 tumour after (a) 0 hours, 24 hours, (c) 48 hours, (d) 72 hours, and (e) 96 hours. After one day (b) there is very little change in the tumour composition as cells remain almost purely green, suggesting no change in their cadherin ratings from zero. After two days (c) small, faintly mesenchymal clumps form on the left and right of the tumour. Upon completing day three (d), early stages of clump formation begin to appear and multiple patches of red mesenchymal cells show scattering throughout the tumour. After the final day (e), multiple clumps of mesenchymal cells have advanced throughout the cadherin rating and progressed rapidly through the stages of EMT.

**Figure 8:**
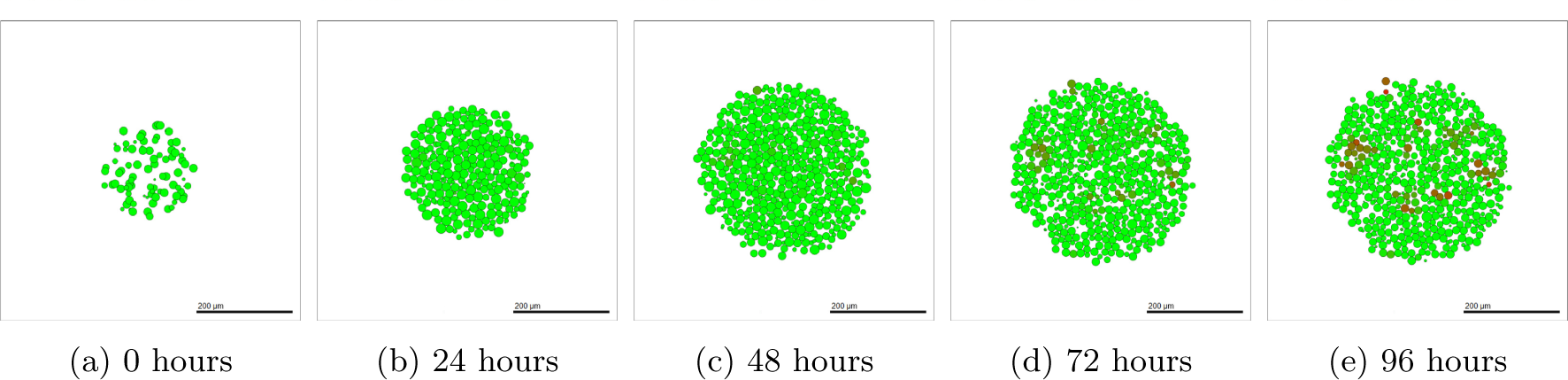
Simulated OVCAR-3 tumour at different time points using default parameter values. Small red mesenchymal clumps begin to form towards the later stages of the simulation.

We observe that despite the small red clumps, the majority of cells remain epithelial with a low cadherin rating. While the volume of these clumps appear negligible in comparison to the volume of the tumour, these clusters of mesenchymal cells cannot be overlooked. Upon breaking away, these cells can relocate and have a key responsibility in the metastasis of the tumour [88]. These clusters can perform collective migration throughout the body despite the lack of individual cell adhesion [89, 90]. The exact procedure by which this is carried out and made viable is relatively undocumented.

Moreover, each cell in the model has a pressure exerted on it by neighbouring cells, changing the cell cycling rate. We can compare the pressure each cell is under to its proximity to the centre of the tumour taken as the origin of the domain, shown in Figure 9 (a). Qualitatively speaking, there is a general negative correlation between the distance a cell is from the centre of the tumour and the pressure acting upon it. This is in agreement with the results found in the literature [91], allowing us to implement pressure-dependent behaviours to the cells in future model adaptations with confidence. Figure 9 (b) shows the oxygen levels throughout the tumour for each cell. Hypoxia is shown to be achieved in central areas of the tumour in which oxygen levels are lower than observed around the tumour exterior.

**Figure 9:**
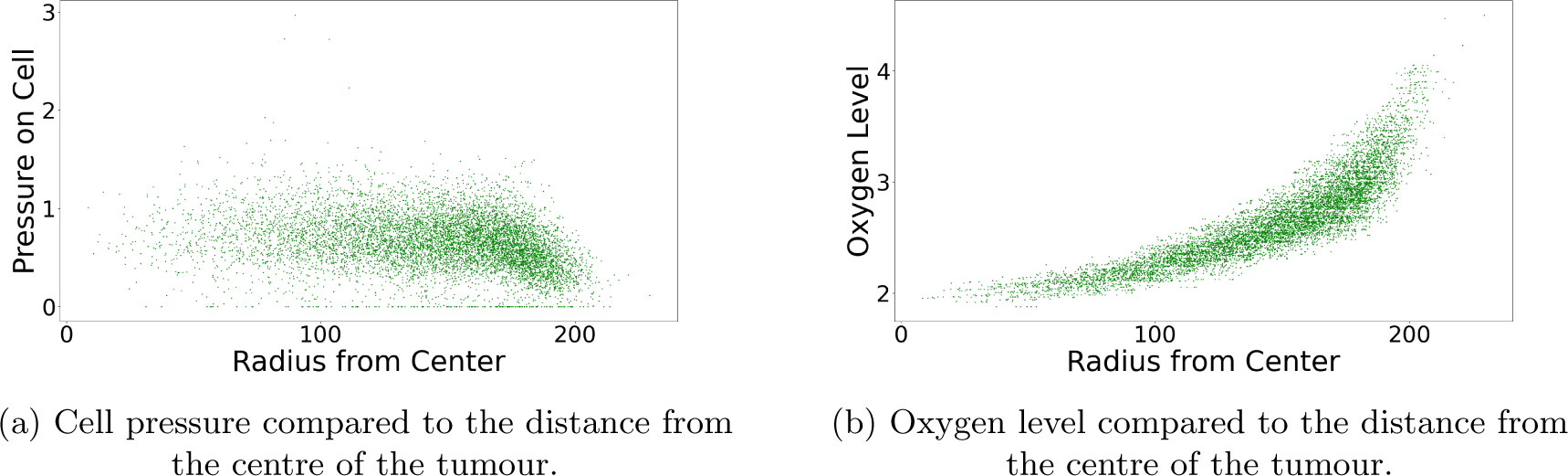
Conditions of cells with respect to the position in the tumour. Each green dot represents a cell at the final time point.

### 3.2 SKOV-3

SKOV-3 tumour simulations give drastically contrasting results to that seen in OVCAR-3 spheroids. Due to the increased jump probability, cells progress through the cadherin ratings faster, as seen after one day in Figure 10 (b). Large areas of the cross-section of the tumour begin to rapidly undergo EMT, creating a blend of epithelial and mesenchymal cells in the neoplasm. After 48 hours of simulated time (c), a clear majority of interior cells have fully undergone EMT and have a cadherin rating of around 13. Occasional green epithelial cells arise around the tumour periphery where oxygen has increased above the threshold value. After 72 and 96 hours in (d) and (e) respectively, the epithelial shell begins to form and creates a solid coating around the primarily mesenchymal tumour. The SKOV-3 tumours show a close agreement to experimental observations shown in Figure 1 (b). The outer shell of epithelial cells remains thin, with only mesenchymal cells making up the interior of the tumour.

**Figure 10:**
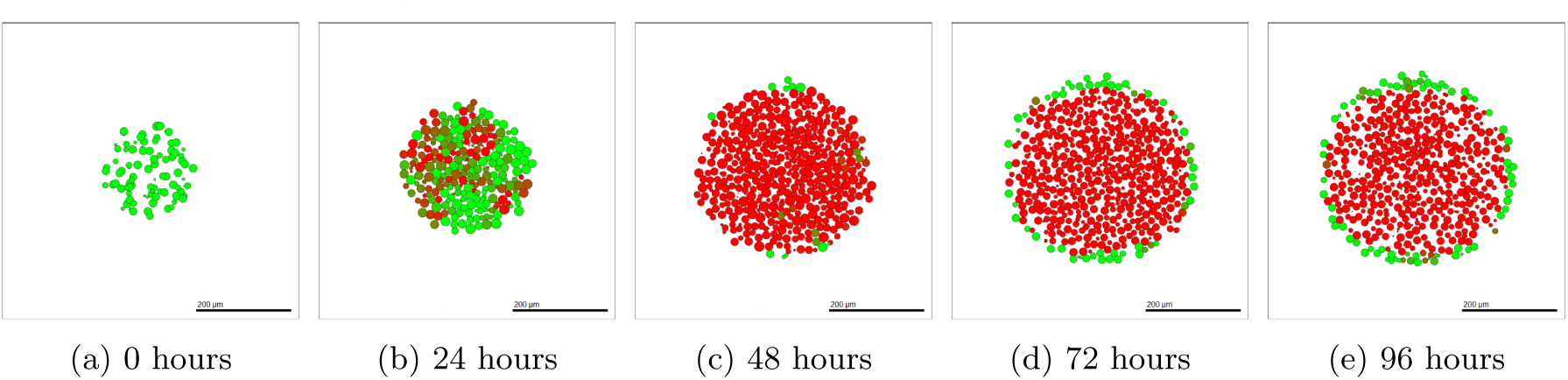
Simulated SKOV-3 tumour at different time points using default parameter values. A red mesenchymal pool quickly forms inside the tumour and a green epithelial layer is created around the periphery.

### 3.3 Model Analysis: Comparison with Experimental Data

Figure 11 shows the proportion of epithelial (E-Cadherin) and mesenchymal (N-Cadherin) cells in the ending population of the tumour. Figures 11 (a) and (b) show the *in-vitro* results while (c) and (d) show the results found *in-silico* using the mathematical model. OVCAR-3 cells finish with a vast majority of epithelial cells in both the experimental and computer simulation results. SKOV-3 cells have a majority of mesenchymal cells, however, due to the outer shell being solely epithelial, SKOV-3 tumours have higher proportions of epithelial cells than OVCAR-3 has mesenchymal. The simulation outputs are in qualitative agreement with the experimental results, with E-Cadherin expression around three times higher in OVCAR-3 tumours than in SKOV-3 tumours. N-Cadherin expression is negligible in OVCAR-3 tumours when compared to expressions in SKOV-3 tumours.

**Figure 11:**
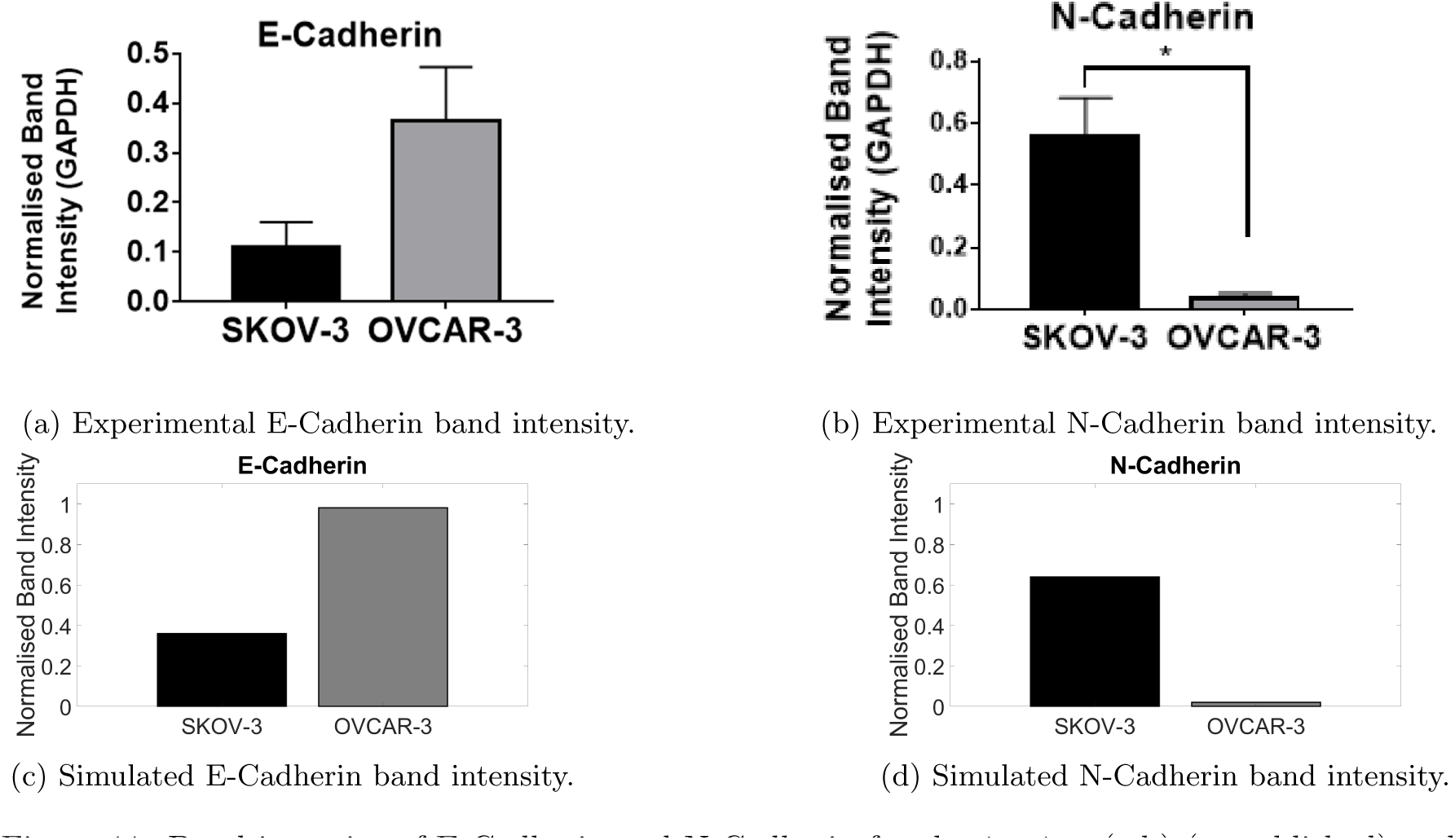
Band intensity of E-Cadherin and N-Cadherin for the *in-vitro* (a,b) (unpublished) and *in-silico* (c,d) experiments after 96 hours.

### 3.4 Sensitivity Analysis

Sensitivity analysis is a crucial part of any mathematical model, quantifying the sensitivity of each parameter on the output of a simulation. By varying these input parameters, we can calculate the variation in the output. Multiple aspects of the output can be investigated, including information on the cell populations or the substrate concentrations. We use a form of Latin Hypercube Sampling to perform global sensitivity analysis. The details of the methods used are described in the Supplementary Material. Using the Pearson Product Correlation (*PCC*) value for different parameters tells us those which have the highest impact on the model output. We compare both how the size of the tumour and the composition of the tumour changes with fluctuations in the parameters used in Equations 1 and 2 for the two cell lines investigated. Generally, all parameters incorporated into these equations have a significant impact on atleast one form of output. Parameters used in Equation 1 tend to have a moderate to high impact on the final simulated tumour size for both OVCAR-3 and SKOV-3 tumours. This is due to changes in the cycling rate having a direct link to the proliferation observed during the simulation time. Parameters used in Equation 2 have less impact on final tumour populations but a much larger impact on the tumour composition. Increasing the parameters in Equation 2 increases the probably of cells undergoing EMT. This impact is more observable in OVCAR-3 tumours where the mesenchymal clumps can be formed throughout the tumour rather than only in the interior. SKOV-3 spheroids generally reach complete EMT in the tumour regardless of small changes in the parameter values used in Equation 2. The results of these sensitivity analyses show that all parameters we have in the generation of the dynamics for the cell cycling rate and jump probability have a large impact and cannot be omitted. This also highlights the importance of finding biologically accurate parameter values, as small perturbations in their value can significantly affect the simulation results.

## 4 Modelling Mesenchymal to Epithelial Transition

It has been observed that mesenchymal cells which have undergone EMT and have relocated to a secondary location may undergo MET at this new location [92]. This allows the cell to reobtain the epithelial phenotype and behaviours previously exhibited to encourage stability and enhanced proliferative abilities [59]. MET has been observed in OVCAR-3 cell lines in which partial EMT has been completed [93]. SKOV-3 cells also show capabilities of showing molecular changes consistent with MET, transitioning from elongated to cuboidal shapes[94]. Here, we explore the effects of MET by including a probability in which cells can jump down in cadherin rating, *q*. Hyperoxic conditions have been found to increase the conversion of EMT into MET within cells [95]. Biological observations also suggest hybrid cells possess the highest cell plasticity on the epithelial-mesenchymal scale [84]. This suggests MET is more likely to occur in hybrid cells than the stable mesenchymal cells. To incorporate these dynamics, we denote two new variables, *bp^∗^* defined as 1 *− bp^∗^* and *oi^∗^* defined as 1 *− oi^∗^*. These represent the contributions of higher oxygen levels and lower cell cadherin rating to an increased probability of MET on each iteration. In terms of probability, these can be seen as the compliments of *bp^∗^* and *oi^∗^* respectively. We then use a similar method to that done in section 2.3.4, as shown in Table 4.

**Table 4:**
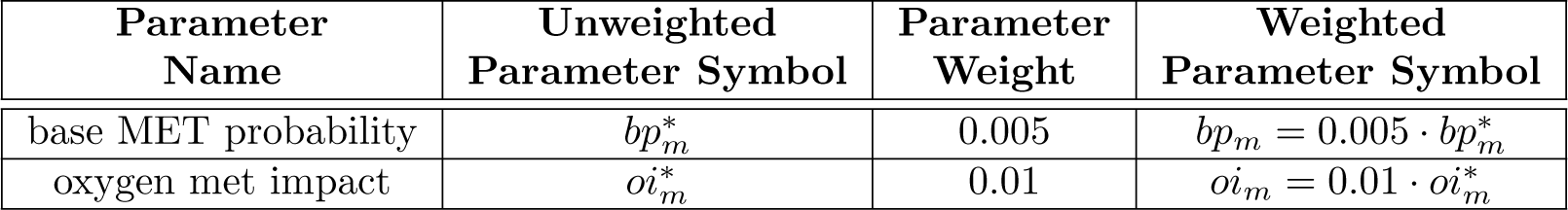
Weightings for parameters used in generating the rates of MET for mesenchymal cells. Only the current cadherin rating and the oxygen levels around the cell change the rate at which a cell can move down the cadherin level.

| Parameter Name | Unweighted Parameter Symbol | Parameter Weight | Weighted Parameter Symbol |
| --- | --- | --- | --- |
| base MET probability | $bp_m^*$ | 0.005 | $bp_m = 0.005 \cdot bp_m^*$ |
| oxygen met impact | $oi_m^*$ | 0.01 | $oi_m = 0.01 \cdot oi_m^*$ |

Using these variables, we can calculate the probability for a cell to jump down in rating during each iteration using Equation 3. Cells are assigned a higher probability when their cadherin rating is lower and the cell is in higher concentrations of oxygen. We use the same “jump down” probabilities for the two cell lines for simplicity.

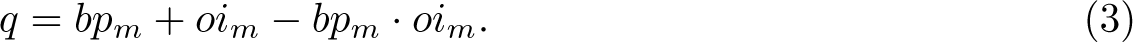

### 4.1 Impact of Initial Conditions

To investigate the effects of heterogeneous cellular composition, a “hybrid” cell classification is now included into the model. Instead of only including epithelial and mesenchymal cells, the population is divided into three subgroups. Cells with a rating of four or less are now classified as epithelial, 5-8 inclusive as hybrid cells, and nine or more as mesenchymal. The tumour is initialised with each cell type to investigate the impact of the starting tumour population type. Epithelial cells are initialised with a rating of two, hybrid with a rating of seven, and mesenchymal with a rating of thirteen. Epithelial cells are no longer initialised with a rating of zero since MET is now included. Starting epithelial cells with a rating over zero allows immediate MET to occur since the cells have been initialised with partial EMT completed.

### 4.2 OVCAR-3

#### Epithelial

OVCAR-3 tumours initialised with epithelial cells no longer obtain the defined isolated clumps of mesenchymal cells observed in Figure 8 during the simulation. Instead, the mesenchymal and hybrid cells develop evenly over time throughout the tumour without clear grouping, as shown in Figure 12. The tumour size grows rapidly over time since the composition of the tumour remains mostly epithelial. Epithelial cells have the fastest cycling rate and so the tumour can proliferate at a faster rate than those initialised with hybrid or mesenchymal cells.

**Figure 12:**
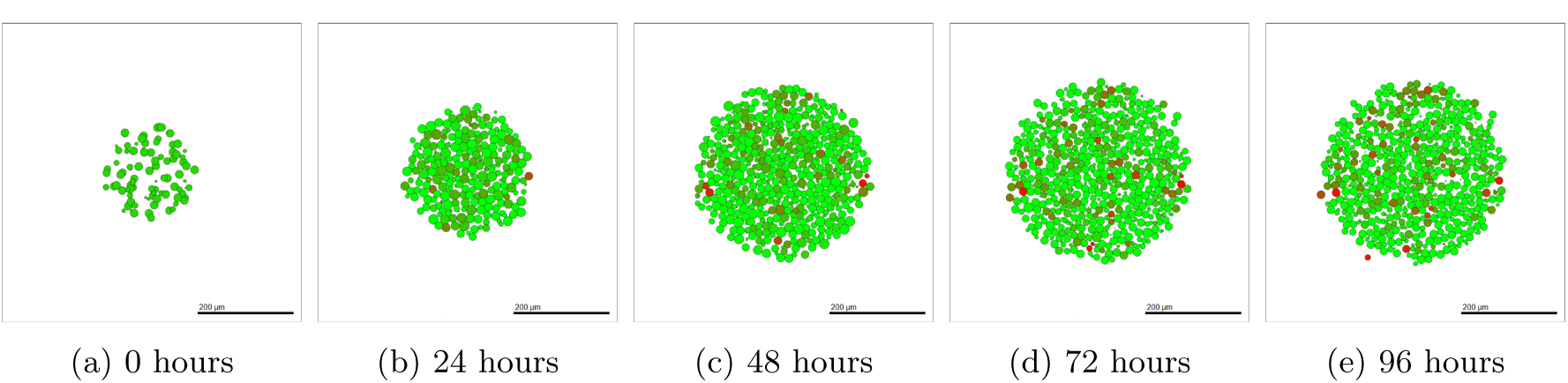
OVCAR-3 tumour over four days of simulated time, initialised with epithelial cells.

The red mesenchymal cells appearing in Figure 12 are spread throughout the entire tumour in what appears to be a generally uniform distribution. The lack of cell adhesion allows mesenchymal cells to break away from the tumour, as seen in Figure 12 (e). The escaping mesenchymal cells in a future model could relocate to a secondary location, representing metastasis in a patient.

#### Hybrid

OVCAR-3 tumours initialised with hybrid cells result in a complete mix of epithelial, hybrid, and mesenchymal cells. No ordering or formation of clumps is visible despite the bystander effect present, as shown in Figure 13. Populations sizes of the cell types appear generally similar with no significant majority of epithelial, mesenchymal, and hybrid cells present.

**Figure 13:**
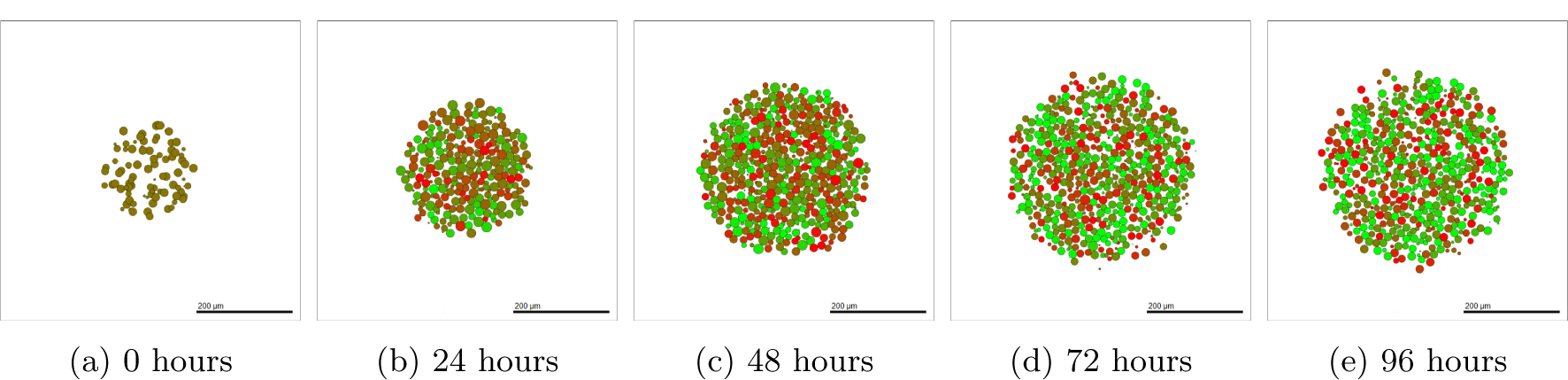
OVCAR-3 tumour over four days of simulated time, initialised with hybrid cells.

The tumour initialised with hybrid cells shows very high plasticity after only 24 hours of simulated time, as seen in Figure 13 (b). Following one day of simulated time there is a significant population of green and red cells at the extremities of the epithelial-mesenchymal scale. This is unlike other initial conditions in which immediate changes in the tumour appearance are not observed as clearly. This suggests that hybrid cells in the model have the lowest stability and can fluctuate between states more readily than the more stable epithelial or mesenchymal cell type.

#### Mesenchymal

When OVCAR-3 tumours are initialised with mesenchymal cells, the majority of the cells remain mesenchymal throughout the four day simulation. After the final simulation output, a scattering of epithelial cells is present throughout the tumour with no immediate observable patterning, as shown in Figure 14.

**Figure 14:**
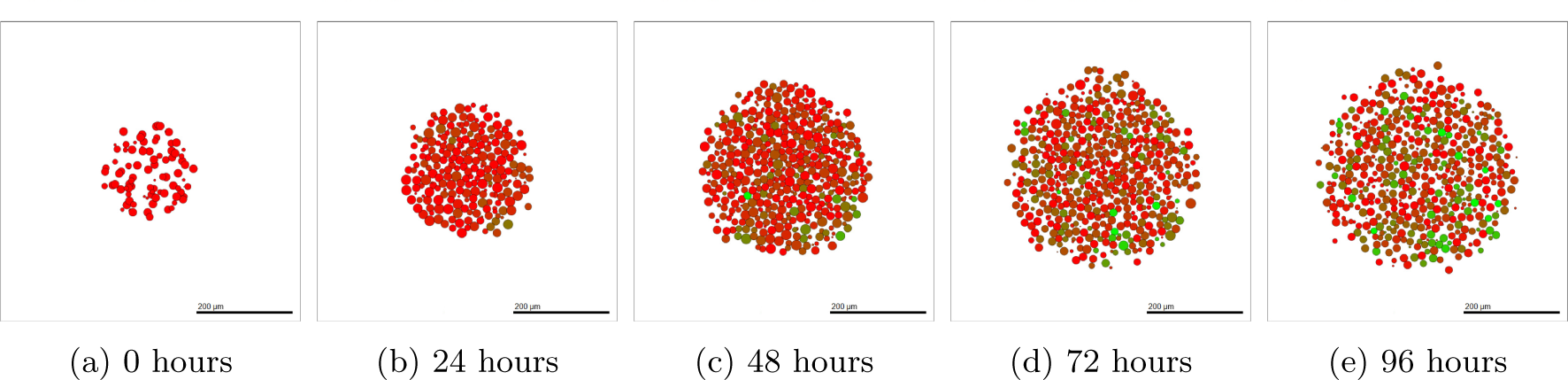
OVCAR-3 tumour over 4 days of simulated time, initialised with mesenchymal cells.

The OVCAR-3 tumour remains mostly mesenchymal over time, with a small number of epithelial cells appearing in the final tumour in Figure 14 (c) and (d). A number of cells can be seen escaping the main tumour clump and moving freely out into the domain. The tumour appears less grouped together with more empty space between the cells than seen in Figures 12 and 13. This is due to the high mesenchymal population decreasing the adhesion strengths within the tumour and the extra cell motility creating a less rigid tumour structure.

#### 4.2.1 Temporal Dynamics

Here, we study the temporal evolution of cell populations to understand the dynamics and cellular transitions as shown in Figure 15. This is compared with experimental data obtained by Ruscetti et al [96] for prostate cancers, obtained using the PKV cell line. The experiments by Ruscetti et al [96] shows the epithelial-mesenchymal plasticity in the PKV cell line cultured *in-vitro*. Population fractions of each phenotype are tracked over the course of 14 days, obtaining the temporal dynamics of the tumour. The PKV cells were isolated using fluorescence-activated cell sorting, identifying epithelial, hybrid, and mesenchymal-like cells during the *in-vitro* experiment. Cells undergoing EMT express green fluorescent protein, suggesting higher rates of transition to a mesenchymal phenotype.

**Figure 15:**
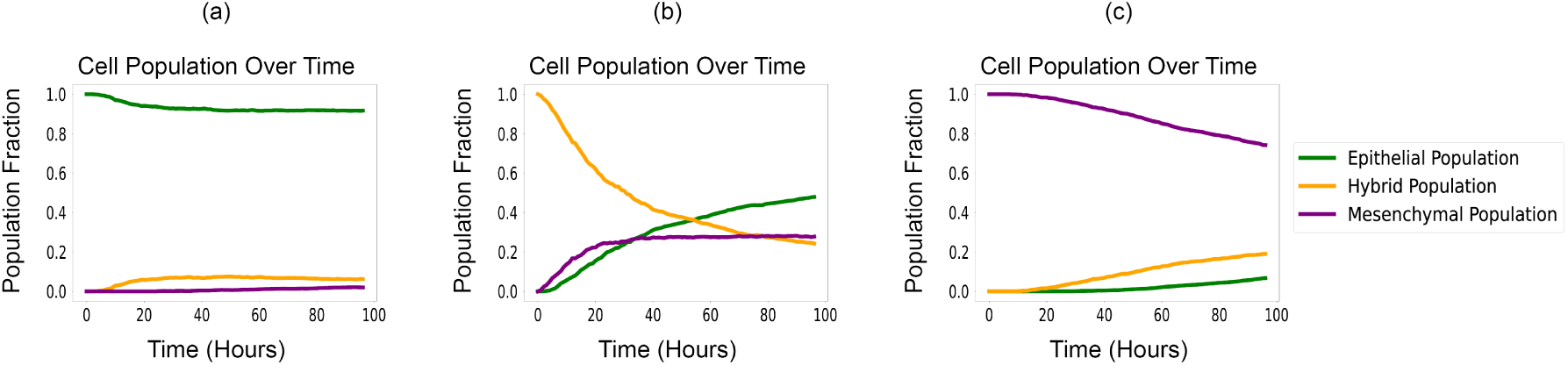
Epithelial vs hybrid vs mesenchymal OVCAR-3 cell populations found *in-silico* including MET.

Experimental data presented in this model uses EMT marker profiles for 2 and 3D cultured SKOV-3 and OVCAR-3 populations characterised in monoculture and adipocyte co-culture. Differences between profiles of 2- and 3D cultured populations were determined via RT-qPCR. For fluorescence analysis of tumour spheroids, immunostaining was carried out. Spheroids were blocked overnight, followed by incubation with primary antibodies for E-Cadherin and N-Cadherin for 48H and secondary antibodies containing fluorophores Alexa 594 (Intrivogen) and Alexa 488 (Intrivogen). Prior to visualisation, spheroids were co-stained with Hoechst and transferred to microscope slides.

The comparisons between SKOV-3 and OVCAR-3 tumours in previous sections highlight the necessity for identifying potential differences in cell line characteristics. Despite this, different cell lines may still possess similar general trends and provide tentative insights into the predicted behaviour of others for model validation. The qualitative trends between the PKV cell line experimental results found *in-vitro* by Ruscetti et al [96] and the OVCAR-3 cell line model results found here *in-silico* are in close agreement. Simulations initialised with epithelial cells remain mostly epithelial, with only small populations of hybrid and mesenchymal cells forming over time, as shown in Figure 15 (a). In Figure 15 (b), hybrid cells can go either way along the cadherin scale, with a significant population of epithelial, hybrid, and mesenchymal cells all present at the final time. Simulations initialised with mesenchymal cells remain mostly mesenchymal, with a non-negligible amount of hybrid cells forming over time and a very small population of epithelial cells.

Figure 16 confirms the importance of initial conditions on the tumour development. Figure 16 (a) and (c) show the stability in tumours initialised with epithelial or mesenchymal cells. While small changes occur in the tumour composition, the majority of cells in these scenarios remain the same type as those they were initialised with. A tumour initialised with hybrid cells (b) shows increased plasticity and fluctuation in the cell types. Hybrid cells end the experiment with the lowest population, with no more than 50% of the tumour made up by a cell of a certain type.

**Figure 16:**
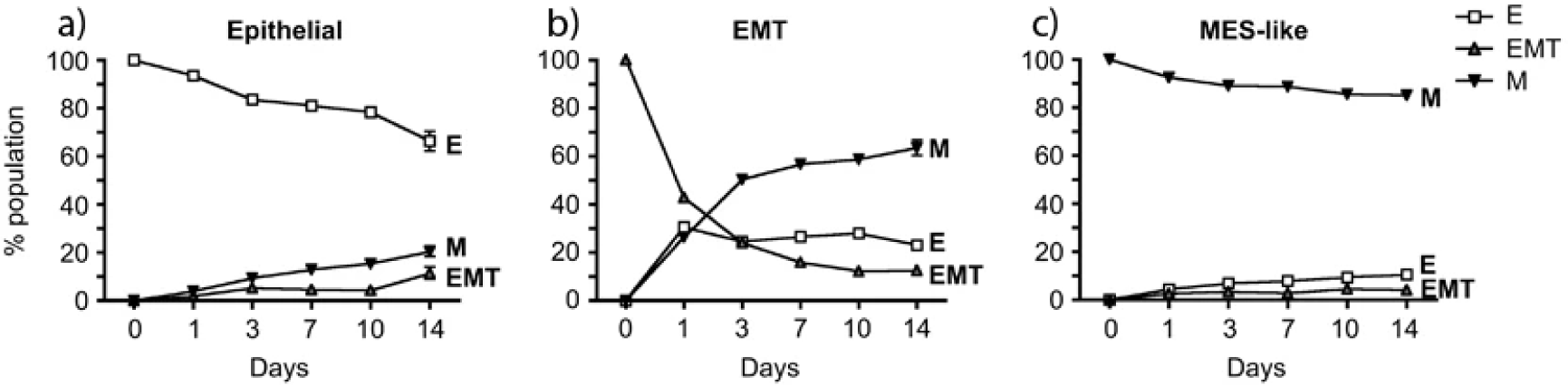
Epithelial vs Hybrid vs Mesenchymal cells populations found *in-vitro* by Ruscetti et al [96].

### 4.3 SKOV-3

We can perform a similar experiment using SKOV-3 tumours. Since SKOV-3 cells are assumed to progress through EMT more rapidly than OVCAR-3 cells, the tumour progresses to the steady state of mesenchymal cells in the interior with a shell of epithelial cells around the exterior very rapidly. This leads to less dependence on the initial condition of the tumour cell type. Figure 17 shows the progression of the tumour over four days when initialised with epithelial cells. After 24 hours of the simulation, the majority of the cells become mesenchymal and have undergone EMT, as shown in Figure 17 (b). Few epithelial cells remain due to sufficient oxygen levels preventing EMT from occurring, with the vast majority of interior cells undergoing complete EMT within the first day. Midway through the simulation, as shown in Figure 17 (c), all cells other than those initiating the formation of the epithelial cells along the periphery undergo complete EMT. These interior cells remain mesenchymal for the remainder of the simulation.

**Figure 17:**
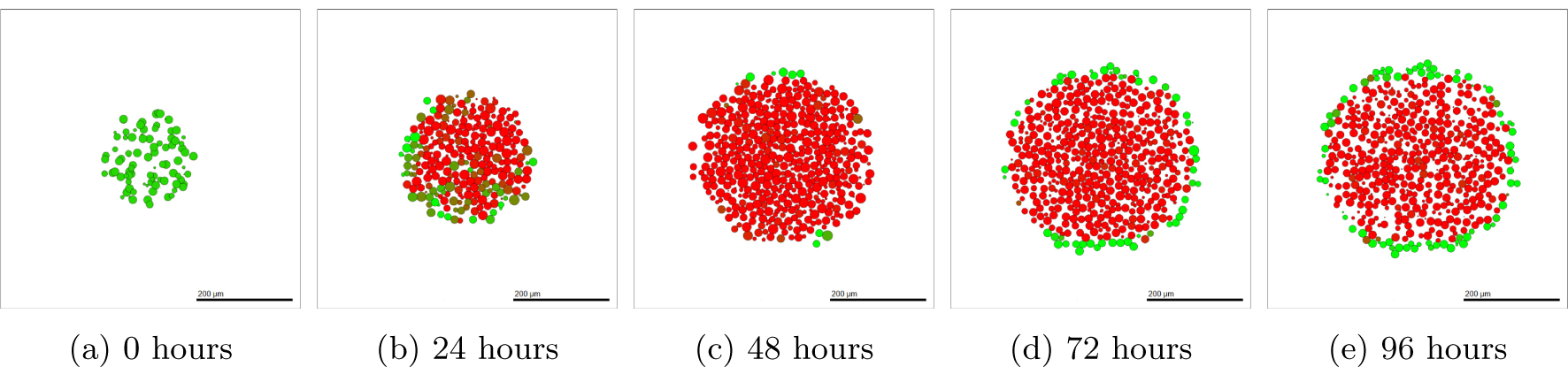
SKOV-3 tumour over 4 days of simulated time, initialised with epithelial cells.

Figure 18 shows the populations of each cell type over time. In all initial conditions, mesenchymal cells rapidly become the main cell type in the SKOV-3 tumour. The epithelial cell population fraction gradually increases in Figures 18 (b) and (c). This is due the tumour periphery expanding outwards and becoming more oxygenated due to to the Dirichlet conditions from the domain boundary. This increased oxygen allows the threshold to be reached deeper into the tumour and the thickness of the epithelial shell to be increased. Cells mostly remain divided into either mesenchymal or epithelial, with very few hybrid cells appearing throughout the tumour. This agrees with the biological observations in the experiments shown in Figure 1 (b), where there is a clear division between the red pool of mesenchymal cells in the tumour core and green epithelial cells around the edge. Regardless of the initial conditions, ending population sizes for each type of SKOV-3 cell remain relatively consistent due to the fast rate at which tumour stability is reached compared to OVCAR-3 tumours.

**Figure 18:**
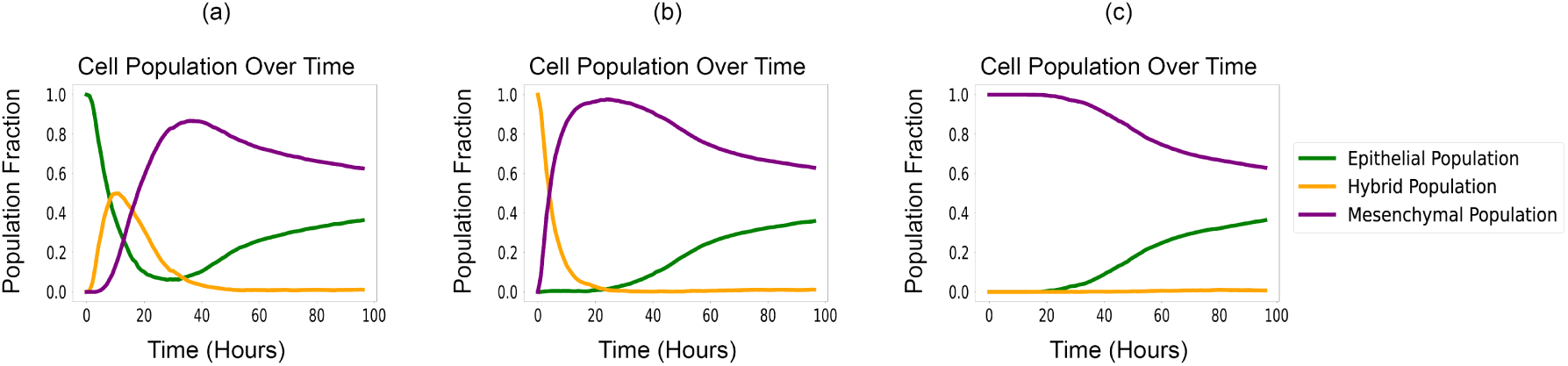
Epithelial vs hybrid vs mesenchymal SKOV-3 cell populations found *in-silico* including MET.

## 5 Conclusions and Discussions

The role of EMT has been shown to have a significant impact on OVCAR-3 and SKOV-3 tumours over time. While on the surface it may appear EMT is a binary process in which a switch is simply turned on, the complex dynamics in the background lead to a microenvironment-dependent, heterogeneous tumour layout. Few models created previously have investigated this continuous perspective of EMT in such a spatially dependent, heterogeneous approach. OVCAR-3 and SKOV-3 cells use similar rules in the mathematical model, with identical Hill functions used in the generation of parameters required for the EMT and cycling rates equations. By varying only the factor at which these parameters are larger in SKOV-3 cells than OVCAR-3 cells and incorporating an oxygen threshold in which SKOV-3 cells become epithelial, the biological observations seen *in-vitro* can be accurately recreated. These changes are sufficient to induce the drastic differences seen between the cell lines experimentally.

The bystander effect is proven to generate results similar to those found experimentally. By using biological observations for calibration, we test our model against other data sets, observed in Figure 16. The results are also compared with models previously designed in similar settings, as seen in Figures 15 and 16. The results show that the developed mathematical model can qualitatively predict the biological observations, indicating the usefulness of the model in exploring processes involved in EMT, MET and potentially to study responses to the therapy.

Sensitivity analysis on the model demonstrates that the outputs of interest are sensitive to all parameters involved in generating the rate at which a cell cycles (Equation 1) and undergoes EMT (Equation 2). SKOV-3 tumours have the unexpected behaviour of the mesenchymal fraction appearing independent of the parameter values used in the jump probability Equation 2. Instead the fraction is dependent on the parameters used in the cell cycling rate in Equation 1. This is due to the fact that oxygen has Dirichlet conditions on the boundary edges, meaning cells able to get closer to the edges are in higher concentrations of oxygen. Higher cycling rates allow the tumour to expand and reach the boundary edges faster, allowing oxygen to reach further into the tumour surface and creating a thicker shell of epithelial cells around the tumour periphery.

The inclusion of MET in Section 4 highlights the importance of including all relevant processes on the cells studied. Including MET restricts the ability of the chemical signal to create mesenchymal clumps in OVCAR-3 tumours via the bystander effect. The tumours without MET ability are more representative of pre-metastatic tumours rather than those which have relocated into a secondary location. The detachable mesenchymal clumps and lack of clear overall structure to the tumour generally make OVCAR-3 tumours more harmful than SKOV-3 tumours. When including the process of MET alongside EMT, differences in the final tumour layouts appear. The clumps seen in OVCAR-3 tumours are less defined and more scattered throughout the neoplasm, while SKOV-3 tumours remain possessing a similar green epithelial shell around the pool of central red mesenchymal cells. By changing initial conditions, we can see the importance of how we set the tumour composition used when starting a simulation. The bystander effect induces a chain reaction, encouraging cells in the proximity of surrounding mesenchymal cells to undergo EMT at an increased rate and become mesenchymal themselves. In SKOV-3 tumours the jump probability is large enough to reach stability within four days as EMT can be achieved in a shorter period of time than in OVCAR-3 tumours. This highlights the need for accurate diagnosis when a patient is seen with ovarian cancer, in both the tumour size and composition. We see a significant difference between the simulations involving only EMT and those also including MET.

The developed multiscale model shows a reasonable level of quantitative and qualitative agreement with experimental data [96]. Despite both different methods of fluorescence analysis performed between the varying cell lines, the general trends found between Ruscetti et al [96] and the mathematical model described above show similar qualitative results. While Ruscetti et al [96] uses a different cancer type (prostate) to that used for our model, this agreement allows us to have a certain level of confidence when extrapolating the model and parameter values beyond those in scenarios investigated experimentally. Doing this creates an opportunity to explore heterogeneous tumour cells with intra-tumour and inter-tumour variabilities such as a specific size, cell line, and mesenchymal composition. This potentially allows one to develop a digital twin and test different treatment dosages and administrative intervals to find the best outcome for each specific patient. Without the aid of mathematical models, creating digital twins in a laboratory setting and investigating multiple different conditions is virtually impossible. These digital twins can conclude which treatment protocol is optimal, thus improving ovarian cancer survival rates and reducing the number of deaths it causes each year.

## Acknowledgments

SO was supported by EPSRC Maths DTP 2021/22 Swansea University [Grant EP/W523963/1]. This research was supported in part by the International Centre for Theoretical Sciences (ICTS) for participating in the program - Theoretical approaches in cancer progression and treatment (code: ICTS/MATHONCO2024/03). GP, MKJ and SO acknowledge the support provided by Global Wales-IISc Joint Research Partnership Fund.

## Notes

### Competing Interest Statement

The authors have declared no competing interest.

