## Supplementary Materials for "Exploring the role of EMT in Ovarian Cancer Progression: Insights from a multiscale mathematical model"

### 1 Sensitivity Analysis

By utilising an adaptation of Latin Hypercube Sampling, global sensitivity analysis can be performed on the model [1]. Analysis is performed on six parameters: $ci_c$ (cadherin impact), $pi_c$ (pressure impact), and $oi_c$ (oxygen impact) used to calculate the cell cycling rate in Equation 1, as well as $bp_e$ (base probability), $oi_e$ (oxygen impact), and $si_e$ (signal impact) used to calculate the EMT jump probability in Equation 2. The values of each parameter is ranged by $\pm 20\%$ using intervals of 2%, leading to 21 different values for each parameter. Each of these 21 values for each parameter is then assigned to one of the 21 simulations we run, resulting in a unique value for each parameter across the simulations. Two output variables from each simulation following 96 hours of simulated time are investigated: the total cell population in the tumour and the fraction of the tumour classified as mesenchymal. This selection is due to the importance of these results in cancer diagnosis. The stage a patient is deemed to be at depends largely upon the size and metastatic ability of the tumour. These outputs are then compared with the input parameter values for each of the six parameters of interest, with the Pearson Product Correlation ($PCC$) value calculated to find the nature and magnitude of the correlation between the $i^{th}$ input $x_i$ and output $y_i$ (See Equation 1) [2]. The value of this $PCC$ variable shows if the correlation between the input parameter and the simulation output is weak, moderate, or strong (See Table 1).

$$PCC = \frac{\sum_{i=1}^{n}\left((x_i - \bar{x})(y_i - \bar{y})\right)}{\sqrt{\sum_{i=1}^{n}(x_i - \bar{x})^2 (y_i - \bar{y})^2}}. \tag{1}$$

| **Coefficient Magnitude** | **Strength of Correlation** |
|---|---|
| 0 | No Correlation |
| Up to 0.4 | Weak Correlation |
| 0.4 up to 0.7 | Moderate Correlation |
| Over 0.7 | Strong Correlation |
| 1 | Perfect Correlation |

Table 1: Evaluations of different values for the Pearson Product Coefficient. Positive (/negative) values suggest a likely positive (/negative) correlation [3]. Stronger correlations between the input and output result in higher magnitudes of the coefficient.

### 1.1 Sensitivity Analysis: Total OVCAR-3 Population

Figure 1 shows the relationship between the key model parameters highlighted previously and the total population of OVCAR-3 cells after 96 hours of simulated time. The impact of the current cadherin rating (a) and pressure the cell is under (c) on the cell cycling rate both have a moderate impact on the final population size. The three parameters used in calculating the jump probability in Equation 2 all have a weak correlation to the total population of OVCAR-3 cells. This is to be expected since these parameters have no direct link to the cycling rate of the cell, instead only affecting the cadherin rating and rate of EMT in the tumour.

### 1.2 Sensitivity Analysis: Mesenchymal Fraction of OVCAR-3 Tumours

Figure 2 shows the relationship between the six parameters and the fraction of the total tumour population considered mesenchymal after 96 hours of simulated time. The three parameters with a direct link to Equation 2 ((d), (e), and (f)) have a moderately significant impact on the output. These parameters have a clear role in calculating the rates at which a cell can undergo EMT. All other input parameters have a negligible impact, suggesting the model has a reasonable level of stability with respect to the user parameters involved in the cell cycling rate Equation (Equation 1).

The correlation between parameters and the outputs between Sections 1.1 and 1.2 show mostly opposite trends. Positive correlations between parameters in Section 1.1 generally show negative correlations in Section 1.2 and vice versa. This is due to the fact tumours with a higher fraction of mesenchymal cells are made up of slower cycling cells and therefore finish with a lower population than those with higher proportions of epithelial cells. Parameters that encourage EMT to occur therefore generally lead to a reduction in tumour growth, as seen in the PPC values between the sections of sensitivity analysis results.

### 1.3 Sensitivity Analysis: Total SKOV-3 Population

Figure 3 shows the relationship between the same six parameters highlighted previously and the total population of SKOV-3 cells. The impact of the current cadherin rating (a), oxygen concentration in the microenvironment (b), and pressure the cell is under (c) on the cell cycling rate all have a moderate impact on the final population size. This is to be expected since these three parameters all have a direct link to the cycling rate of the cell. All other input parameters have a negligible impact, suggesting the model has a reasonable level of stability in relation to the user parameters.

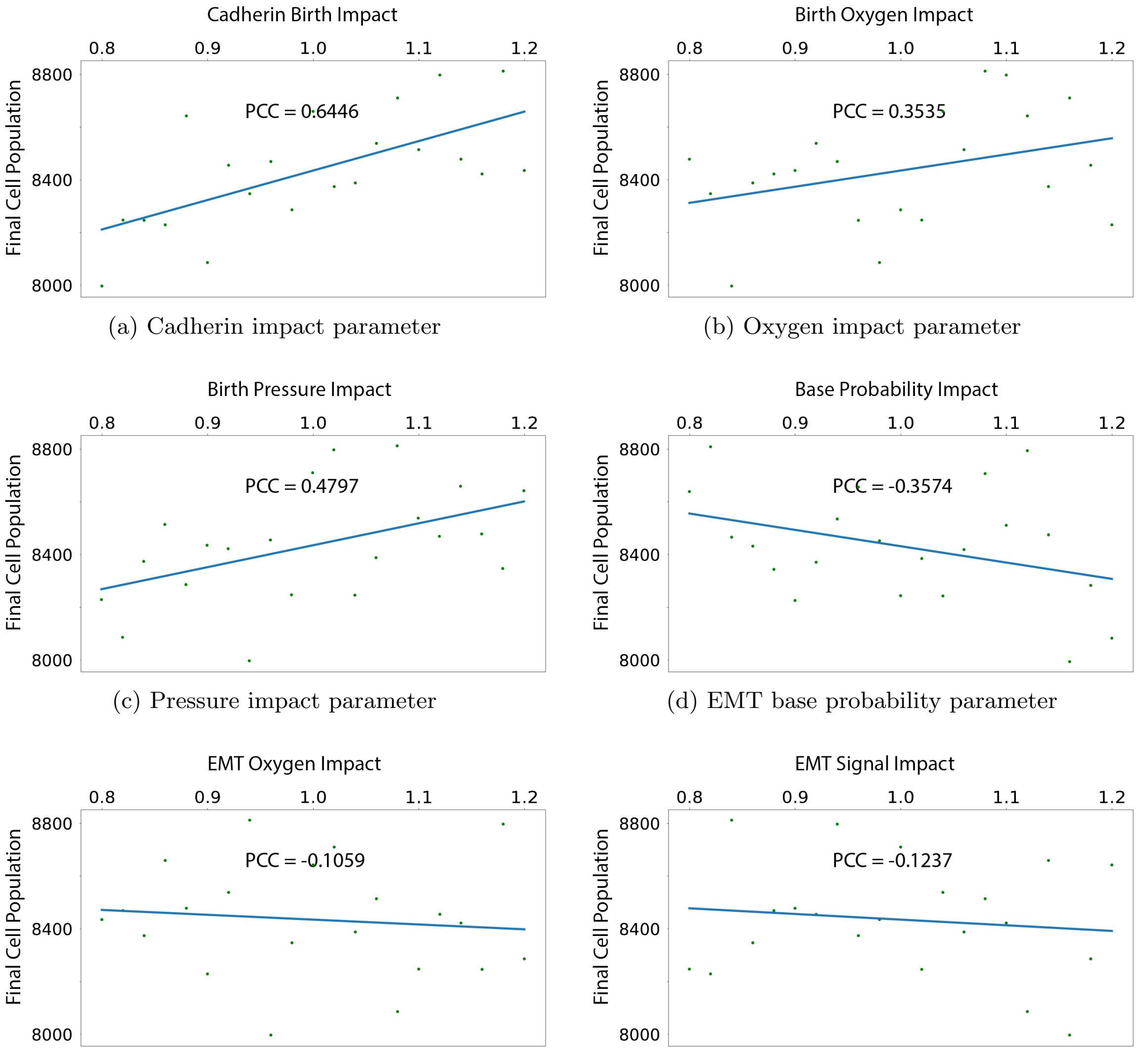


(a) Cadherin impact parameter

(b) Oxygen impact parameter

(c) Pressure impact parameter

(d) EMT base probability parameter

(e) EMT oxygen parameter

(f) EMT signal parameter

Figure 1: Impact of parameter values on the final total cell population in the OVCAR-3 tumour. The fraction of the total tumour cell population classed as mesenchymal is found and compared for various parameter values. PPC is given for each parameter across 21 simulations. Figures (a), (b), and (c) concern the birth rate used in Equation 1, while Figures (d), (e), and (f) concern rates at which EMT can occur within the cells used in Equation 2.

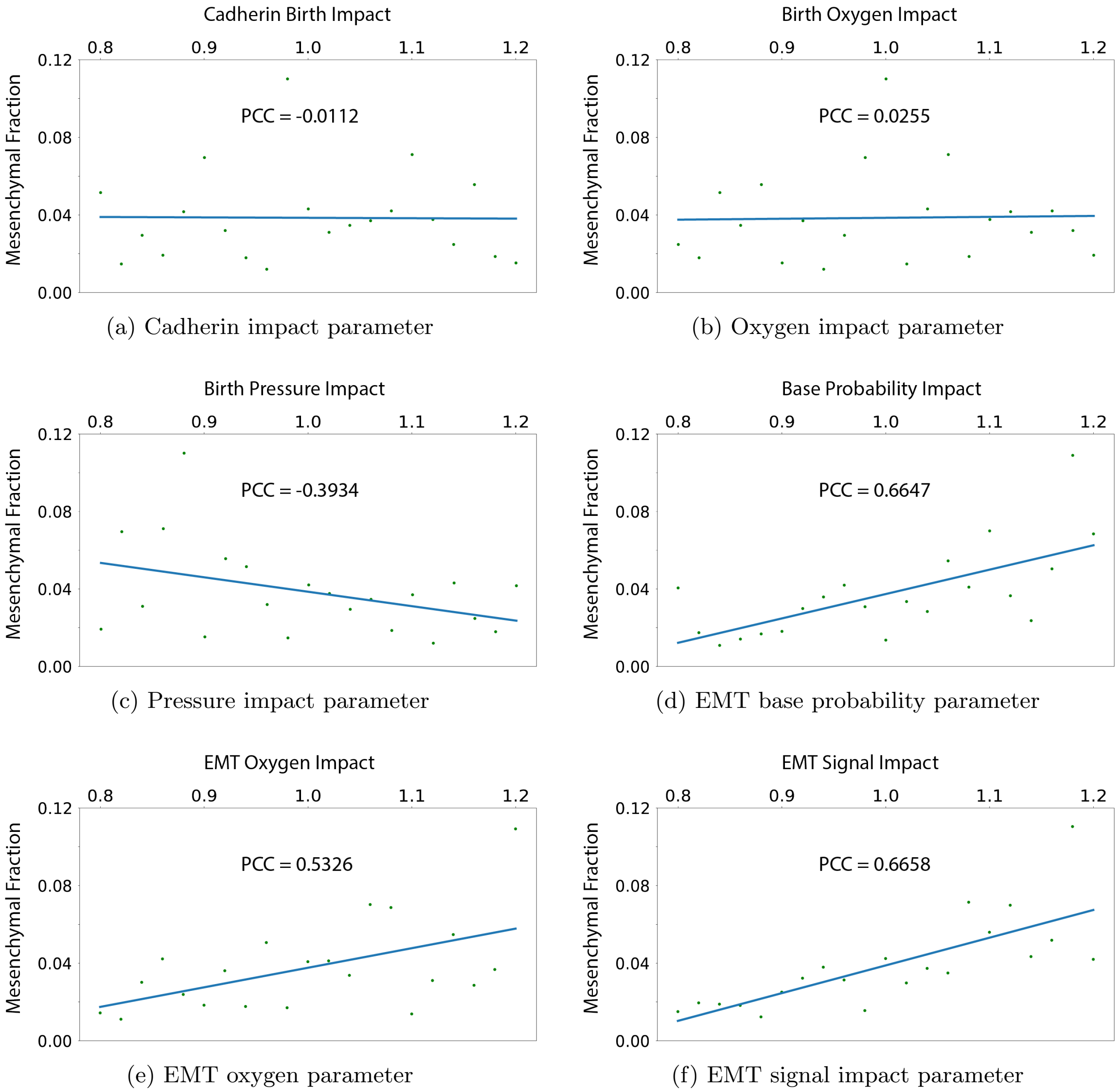


(a) Cadherin impact parameter

(b) Oxygen impact parameter

(c) Pressure impact parameter

(d) EMT base probability parameter

(e) EMT oxygen parameter

(f) EMT signal impact parameter

Figure 2: Impact of parameter values on the final composition of the OVCAR-3 tumour.

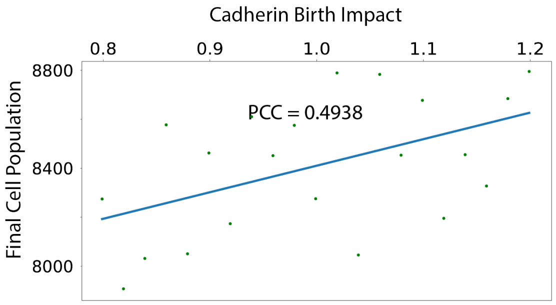


(a) Cadherin impact parameter

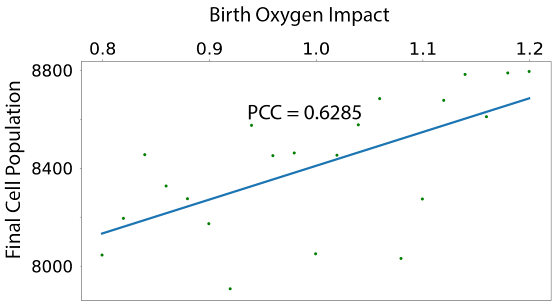


(b) Oxygen impact parameter

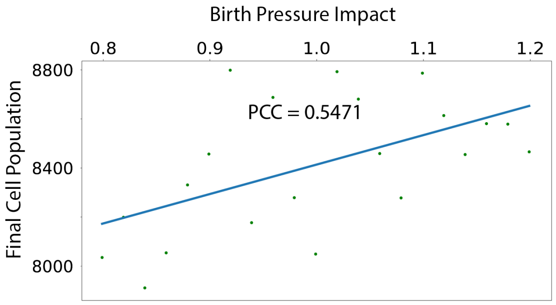


(c) Pressure impact parameter

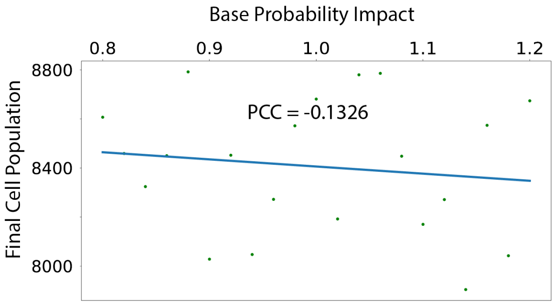


(d) EMT base probability parameter

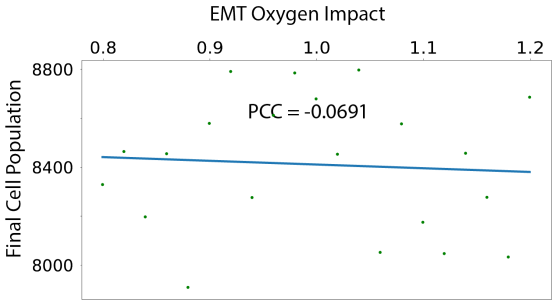


(e) EMT oxygen parameter

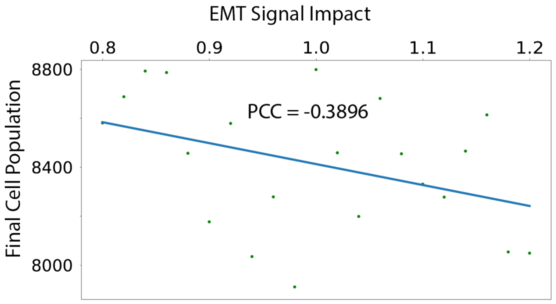


(f) EMT signal parameter

Figure 3: Impact of parameter values on the final total cell population in the SKOV-3 tumour.

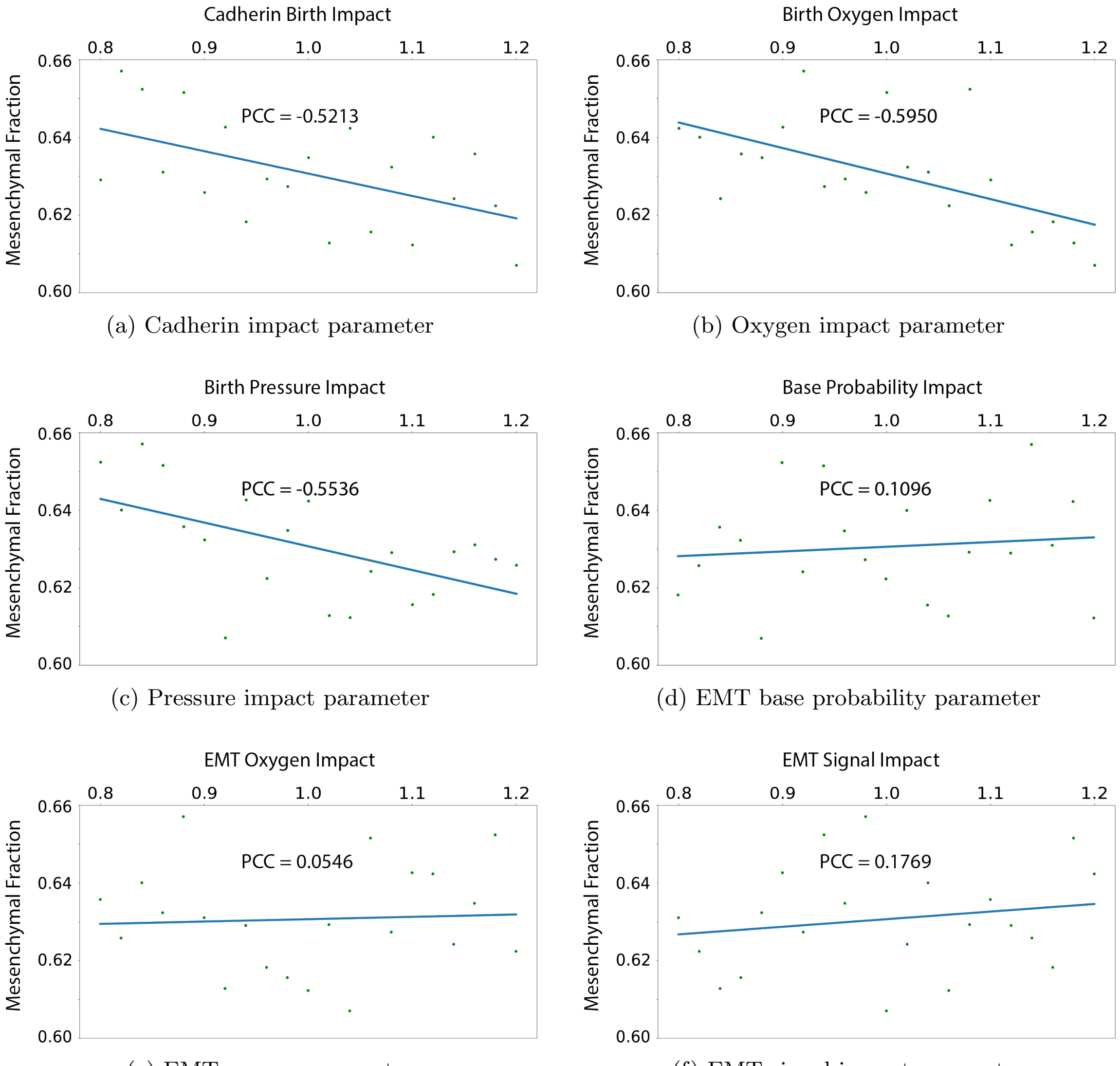


(a) Cadherin impact parameter

(b) Oxygen impact parameter

(c) Pressure impact parameter

(d) EMT base probability parameter

(e) EMT oxygen parameter

(f) EMT signal impact parameter

Figure 4: Impact of parameter values on the final composition of the SKOV-3 tumour.

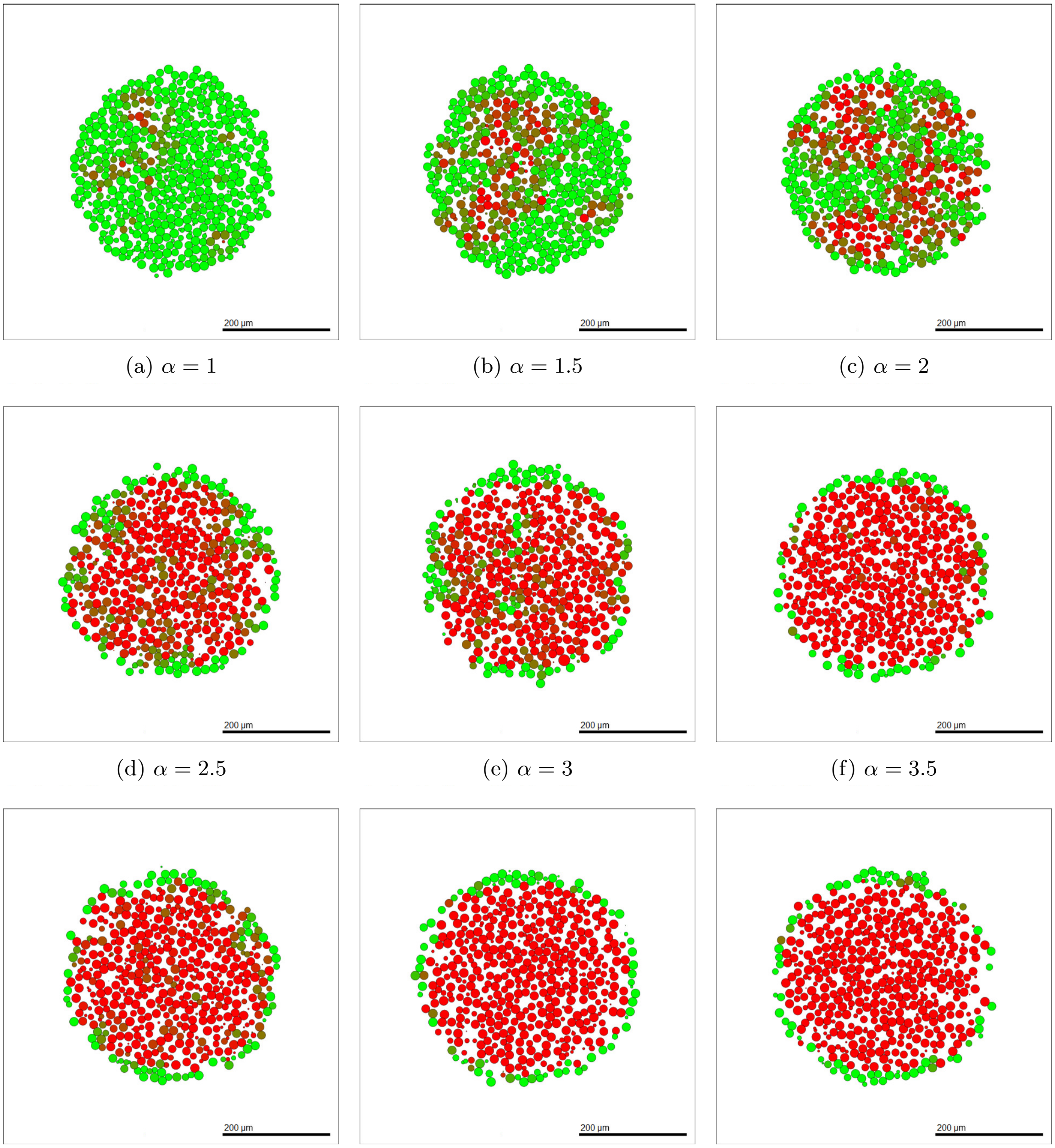


(a) $\alpha = 1$ (b) $\alpha = 1.5$ (c) $\alpha = 2$

(d) $\alpha = 2.5$ (e) $\alpha = 3$ (f) $\alpha = 3.5$

(g) $\alpha = 4$ (h) $\alpha = 4.5$ (i) $\alpha = 5$

Figure 5: Impact of the value of $\alpha$ on the final composition of the SKOV-3 tumour.

#### 1.4 Sensitivity Analysis: Mesenchymal Fraction of SKOV-3 Tumours

Figure 4 shows the relationship between the six parameters and the fraction of the total SKOV-3 tumour population considered mesenchymal after 96 hours of simulated time. Again, the three parameters directly involved in the cell cycling equation (Equation 1) have a moderate impact on the proportions of tumour composition, with all other parameters having a weak correlation. This is less expected since these parameters only directly change the cell cycling rate. Changing this cell cycling rate causes the tumour to expand, causing the shell of epithelial cells around the edge of the tumour to move closer to the domain boundaries. Here the oxygen available to the cells is higher, allowing a thicker shell of epithelial cells, thus decreasing the fraction of the tumour cells deemed to be mesenchymal.

#### 1.5 Sensitivity Analysis: Cell Line Differentiation

Another approach to sensitivity analysis can be to change the difference in weighting terms between Tables 2 and 3. In these tables the weightings of parameters used in the jump probability generation in Equation 2 have five times larger weightings for SKOV-3 cells than OVCAR-3 cells. This creates the clearly distinguishable difference in tumour layouts, with OVCAR-3 possessing disjoint clumps of mesenchymal cells while SKOV-3 tumours have a pool of mesenchymal cells making up the entire interior. We define $\alpha$ to denote the factor at which SKOV-3 cells have a larger weighting than that used for OVCAR-3 cells in Table 2. For example, $\alpha = 5$ in Table 3. We can vary the value of $\alpha$ to explore at which point the tumour shows OVCAR-3 and SKOV-3 characteristics. Figure 5 shows the tumour appearance after 96 hours of simulated time for different values of $\alpha$.

In Figure 5 (b) and (c) where $\alpha = 1.5$ and $\alpha = 2$, the tumour appears to show a hybrid state of OVCAR-3 and SKOV-3. Large clumps of mesenchymal cells are formed, however, due to their increased size these begin to overlap and make up a large proportion of the inside of the tumour. With values of $\alpha$ between two and four in Figures 5 (d), (e) and (f), the pool of mesenchymal cells has formed with only occasional epithelial cells appearing within the tumour interior. This appears closer to the original SKOV-3 tumour appearance. The slow transition between the cell lines across the figures display that the switch in the model between SKOV-3 and OVCAR-3 cells can be continual and non binary, generating cells with characteristics of both cell lines.
